# Emergent neural dynamics and geometry for generalization in a transitive inference task

**DOI:** 10.1101/2022.10.10.511448

**Authors:** Kenneth Kay, Natalie Biderman, Ramin Khajeh, Manuel Beiran, Christopher J. Cueva, Daphna Shohamy, Greg Jensen, Xue-Xin Wei, Vincent P. Ferrera, L.F. Abbott

## Abstract

Relational cognition — the ability to infer relationships that generalize to novel combinations of objects — is fundamental to human and animal intelligence. Despite this importance, it remains unclear how relational cognition is implemented in the brain due in part to a lack of hypotheses and predictions at the levels of collective neural activity and behavior. Here we discovered, analyzed, and experimentally tested neural networks (NNs) that perform transitive inference (TI), a classic relational task (if A > B and B > C, then A > C). We found NNs that (i) generalized perfectly, despite lacking overt transitive structure prior to training, (ii) generalized when the task required working memory (WM), a capacity thought essential to inference in the brain, (iii) emergently expressed behaviors long observed in living subjects, in addition to a novel order-dependent behavior, and (iv) adopted different task solutions yielding alternative behavioral and neural predictions. Further, in a large-scale experiment, we found that human subjects performing WM-based TI showed behavior inconsistent with a class of NNs expressing an intuitive task solution. These findings provide neural insights into a classical relational ability, with wider implications for how the brain realizes relational cognition.

## 1 Introduction

Cognitive faculties such as logical reasoning, mathematics, and language have long been recognized as characteristic of human-level intelligence. Common to these faculties is abstraction: the ability to generalize prior knowledge and experience to novel circumstances. Importantly, abstraction typically entails understanding particular relationships, e.g. “adjacent to”, “same as”, “relevant to”) between items (e.g. stimuli, objects, behaviors, words, variables), which can then be used to infer equivalent relationships between items not previously observed together, i.e. *novel combinations of items*. Such relational inferences – which can also be understood as systematic generalizations to novel compositions of inputs – are the basis of a structured form of knowledge, often termed a “schema,” that is thought to enable humans to generalize in systematic and meaningful ways [1–6], and thus has been posited as essential to advanced cognition.

Intriguingly, landmark work in animals [7–12] indicates that cognition based on relations is more prevalent, and thus possibly more essential, than previously thought. This prevalence is evidenced by the observation that cognitive abilities that entail relational inference – such as navigation [13–15], learning-to-learn [9, 16, 17], and concept/structure learning [11, 12, 18] – are in fact widespread across the animal kingdom. Further, these abilities have been linked to memory systems in the brain – variously termed “relational memory”, “cognitive maps”, “learning sets”, among others – that enable humans and animals alike to make systematic inferences [5, 19–21], generalize across different domains [16, 22, 23], learn rapidly [24–28], and plan and envision new experiences [29–36]. These findings and insights extend the scope of relational inference to a wide range of species and cognitive abilities, and, further, imply that there exists a deep interrelationship between relational inference and memory.

Despite this unifying importance, it remains an open question how relational inference is implemented in neural systems, whether in artificial networks (e.g. those performing linguistic [37–39] or symbolic [40, 41] tasks) or in the brain. Toward answering this question, a fundamental scientific aim is to identify or generate putative neural implementations that can be used to derive empirically testable hypotheses. In particular, hypotheses at the level of behavior and of collective (population-level) neural activity may be crucial given that these levels have proved decisive in clarifying whether and how neural systems in the brain implement various cognitive functions (e.g. vision, movement, timing, decision-making [42–46]). Notably, in both neurobiology and machine intelligence, relational inference is often studied in relatively complex cases (e.g. spatial [16, 21, 24, 47–49] and linguistic [37, 38, 50] knowledge), for which it may be relatively difficult to formulate or generate hypotheses derived from neural implementations, especially at the level of neural populations and of behavior. In this way, studying simpler cases of relational inference may be advantageous or even crucial towards understanding relational inference in the brain. We therefore took a two-part approach intended to yield such hypotheses: first, we stipulated a task paradigm that distills relational inference into a simple yet essential form, and second, we adopted a methodology suited to discover possible population-level and behaviorally relevant neural implementations thereof.

## 2 Transitive inference: a classic cognitive task

We first sought to operationalize relational inference in a task that is (i) reduced to a single abstract relation, and, further, (ii) based on stimuli (e.g. images) that can be presented with temporal precision. We reasoned that each of these properties might be critically important in that (i) reduces complexity, which could ultimately enable discovering neural implementations, and (ii) delimits periods of sensory-driven neural activity, and by extension neural activity underlying inference, thus facilitating subsequent interpretation.

A classic task paradigm capturing these properties is **transitive inference** (**TI**) [10, 51–54], which tests a subject’s ability to use premises A > B and B > C to infer A > C — an instance of use of an abstract rule based on an underlying relation (i.e. choose ‘higher’ items, based on the transitive ‘>’ relation). TI operationalizes a simple yet powerful schema (**Fig. 1a**) that enables generalization from N premises (training cases) to order *N*^2^ probes (test cases) in accordance with the relation-based rule (i.e. choose the ‘higher’ item). In contrast to generalization based on interpolation or extrapolation, TI tests generalization that is expressly compositional and based on an underlying schema (‘schematization’), manifesting behaviorally in the pattern of correct responses in the task (**Fig. 1b**). Transitivity (‘>’) is itself characteristic of many types of relations, and is of fundamental importance in symbolic reasoning – together suggesting that TI is not essentially based on specific stimuli or stimulus features in isolation.

**Figure 1:**
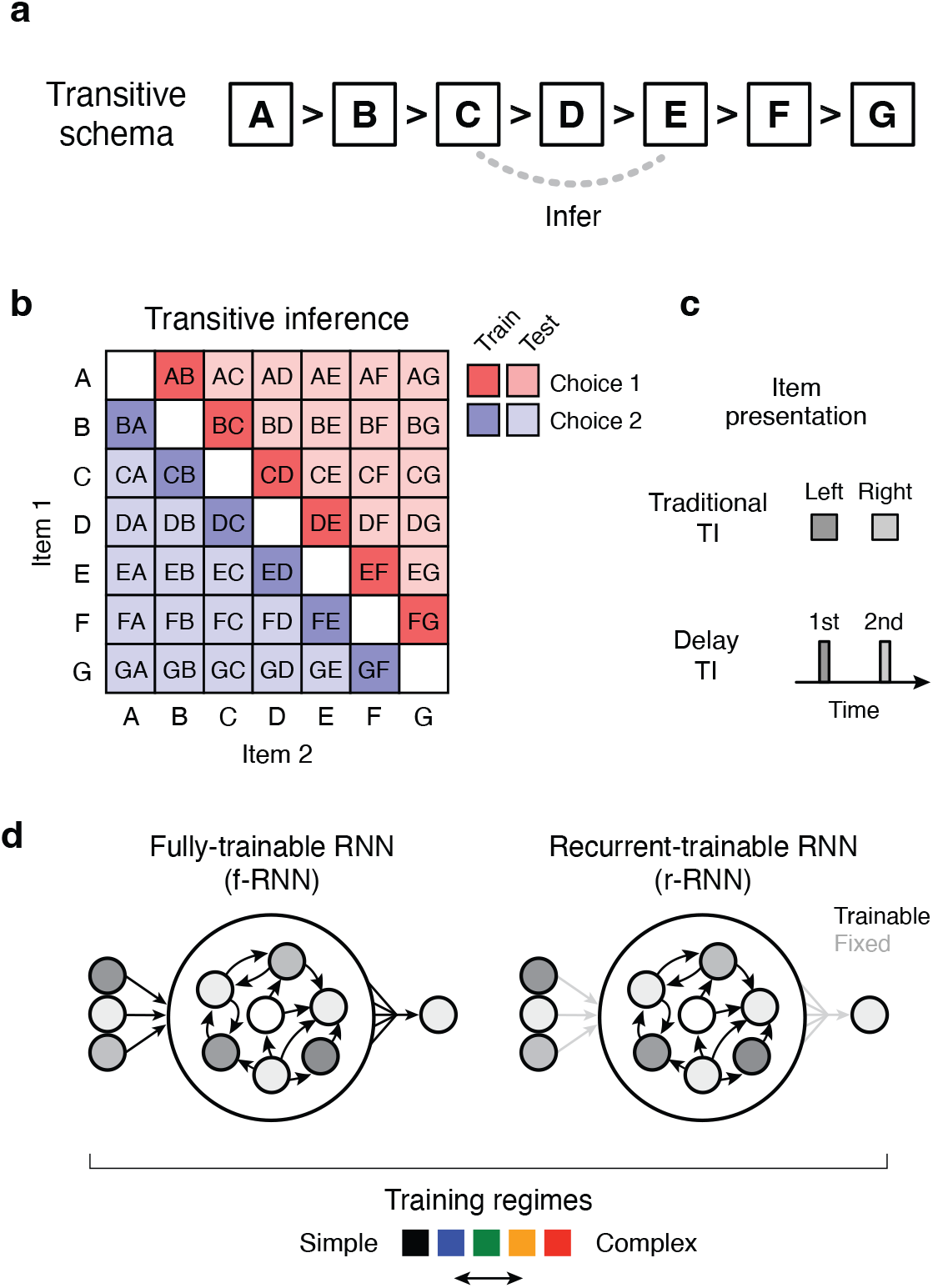
Cognitive task and neural approach. **a**, Diagram of the relational structure (schema) underlying transitive inference (TI). Subjects learn correct responses to premise pairs (A vs. B, B vs. C, C. vs. D, etc.; training trials), and must infer correct responses in previously unobserved (novel) pairs (C vs. E, etc.; testing trials). Correct responses are specified by an abstract rule: for every possible pair, the “higher” item in the schema should be chosen, in accordance with the transitive relation. Importantly, the items in the schema (A, B, etc.) correspond to arbitrary real-world items. **b**, Trial types and their correct responses in 7-item TI. Each trial type consists of a permutation of two items (item 1 and item 2). **c**, Item presentation formats: traditional TI vs. delay TI. In traditional TI, items are presented simultaneously and are chosen on the basis of their presented position (e.g. left vs. right). The present study proposes “delay TI”, in which items are presented with an intervening delay and are chosen on the basis of presentation order (1st vs. 2nd). The delay version explicitly requires working memory (WM), a cognitive capacity thought to be essential for relational inferences in living subjects. **d**, Neural models in the present study. Among neural network (NN) architectures, recurrent neural networks (RNNs) are able to implement WM. Two variants of RNNs were studied: fully-trainable RNNs (f-RNNs), for which both feedforward and recurrent synaptic weights were modifiable in training, and recurrent-trainable RNNs (r-RNNs), for which only recurrent synaptic weights were modifiable in training (feedforward weights randomly generated and fixed; trainable vs. fixed weights diagrammed as black vs. grey arrows, respectively). In conjunction, RNNs were trained with different levels of regularization and initial synaptic strengths (‘training regime’, indicated in later figures by color; parameters in **Table 5**). All networks had 100 recurrent units, which all had the same non-linear activation function (tanh). See Methods for additional details.

At a broader level, TI can be understood as a simplified alternative to other paradigms testing relational inference but involving multiple relations and abstract rules (e.g. linguistic syntax and semantics) and/or task stimuli that are relatively challenging to isolate (e.g. spatial paradigms [16, 21, 24, 47]). Further, in contrast to approaches that focus mainly on lower-level neurobiological phenomena (e.g. neural firing that has abstract correlates [14, 22, 55]), TI requires direct behavioral report of successful inference, thus affording identification of potentially critical relationships between behavior and underlying neural implementations.

Remarkably, though TI is a classic task in behavioral psychology [10, 51, 53, 56, 57] and a cornerstone of symbolic reasoning, the neural basis of TI has remained unclear [53, 58–60], potentially due to a lack of putative neural implementations and testable hypotheses.

## 3 A neural approach to TI

Despite the long history of TI as a cognitive task, investigation of its neural basis in the brain is comparatively recent [52, 61–65]. Prior work implies that an approach seeking to identify biologically accurate neural implementations would benefit from two criteria:

First, an approach should impose **minimal architectural constraints**. TI is observed in an extremely broad range of species, including primates, birds, rodents, and insects [53, 59, 60, 66–68] (possibly the result of convergent evolution [69]), a striking ubiquity that implies that highly specialized neural architecture may not be essential.

Second, an approach should explicitly require memory across time, particularly **working memory (WM)** [70–73]. In living subjects, relational inferences such as TI characteristically rely on memory since subjects must assess relationships between events not experienced simultaneously (e.g. sensory stimuli separated in time) – memory enables such events to be in effect brought together. WM in particular is thought to be essential to relational inference [74–77], not only because WM is generally required in real-world cases of relational inference (e.g. language comprehension, spatial navigation), but also because prior work suggests that relational inference is accomplished in the brain by a neural system that intrinsically supports and/or relies upon WM (e.g. prefrontal cortex, possibly via a process akin to deliberation or reasoning [70,72,78]). Surprisingly, though TI exemplifies relational inference, prior work has only evaluated indirect relationships between TI and WM (either by having separate WM vs. TI tasks [79], or by linking each to a common brain region [61, 80]), with almost no study directly testing WM by imposing an intervening delay between presented items, which we here call “delay TI” (item 1 - delay - item 2; **Fig. 1c** and **Fig. S1a**; see [81] which imposes a WM delay in a probabilistic version of TI). Notably, task paradigms testing TI often have items in separate environmental locations or containers, thus implicitly requiring WM (e.g. [52, 67, 68]).

Importantly, these two criteria are suited for the methodology of generating and analyzing task-trained **recurrent neural networks** (**RNNs**), an approach that has proved successful in discovering neural implementations of other cognitive abilities, and, further, in generating testable predictions at the level of collective neural activity [82–84]. We therefore adopted this approach, and, further, expanded upon it in two ways. First, in conjunction with RNNs, we also assessed whether and how trained models that cannot implement WM, yet have neurally relevant feedforward structure, might transitively generalize. To do so, we investigated two archetypal models: logistic regression (LR [85]) and multi-layer perceptron (MLP [86]) (schematic in **Fig. S2a**), each tested on the “traditional TI” format having no explicit delay (**Fig. 1c**). Second, to identify (where possible) multiple solutions to TI, we investigated two neurobiologically relevant types of RNN variants:

### Learnable connectivity

In NNs and the brain, network architecture is comprised of feedforward and/or recurrent connectivity, the respective roles of which in learning cognitive tasks remain unclear (e.g. [87–89]) – this is indeed the case for TI, a paradigm that requires subjects to learn the premise trials (e.g. A vs. B, B vs. C, etc.). Thus it is unclear whether the neural system implementing TI in the brain relies on learned connectivity that is feedforward, recurrent, or some configuration of both. It is worth emphasizing that this distinction is in fact critical for relational inference tasks such as TI, where the significance of the stimuli within the task (items A, B, C, etc.) has no *a priori* relationship to stimulus features, unlike tasks based on stimulus features known to be encoded in feedforward inputs from upstream brain regions (e.g. tactile frequency [90, 91], visual frequency and orientation [92], object categories [93]). To address this gap in understanding, we investigated RNNs for which feedforward connectivity was either trainable or not trainable, i.e. “fully-trainable” RNNs (f-RNNs) vs. “recurrent-trainable” RNNs (r-RNNs), respectively (**Fig. 1d**).

### Training regime

The accuracy of trained NNs in matching neural systems in the brain has been found to depend on efficiency constraints; these constraints take the form of training penalties that limit excess neural activity (regularization; [48, 89, 94–97]). Further, differences in the initial strength of connectivity – the magnitude of connection weights prior to training [98, 99] – have been found to yield NNs having striking differences in implementation (internal representations and dynamics) that are concomitant to differences in neurobiological accuracy [48, 95, 96, 100, 101]. These findings indicate that it may be critical to investigate both factors (training constraints and initial strength of connectivity) in trained NNs when seeking to identify biologically accurate neural implementations. We thus trained NN variants using parameter sets that systematically varied these factors, referring to each set as a “training regime” (‘simple’ to ‘complex’, following similar terminology distinguishing NN variants in [95], **Table 5**).

## 4 A variety of neural models perform TI

We first sought to determine whether trained RNNs could successfully perform TI. Unlike perceptual tasks or tasks that solely test memory, TI expressly requires generalization to novel combinations of inputs. Such generalization requires some form of additional knowledge regarding the underlying relationship between inputs (the transitive schema, **Fig. 1a**). Thus TI is a task that requires an *a priori* inductive bias – here, for transitivity – the implementation of which is not generally known in relatively unstructured models such as trained RNNs [84, 102, 103].

Mirroring TI as encountered by living subjects (7-item TI with items A to G; each item represented as a random high-dimensional (100-D) input), we trained RNNs (100 recurrent units, tanh nonlinearity) exclusively on premise (training) trials (consisting of ‘adjacent’ item pairs A vs. B, B vs. C, C vs. D, etc.) and evaluated whether RNNs generalized to test trials (B vs. D, etc.; all trial types shown in **Fig. 1b**). Training was conducted using gradient descent optimization and backpropagation-through-time; further, all trials required working memory (WM) to relate items separated by a stimulus-free delay (delay TI; **Fig. 1c** and **Fig. S1**). Response choice was defined by which of two output units (linear readouts corresponding to choice 1 vs. 2) met a fixed activity threshold (85% of maximum value), and response time (RT) was defined as the time this threshold was reached. For an initial assessment of whether the networks could generalize, a simulation of all trial types was performed under noiseless conditions.

We found that trained RNNs often generalized perfectly, i.e. responded correctly to all test trials (example RNNs in **Fig. 2**; summary in **Table 1**; additional RNNs trained on extended and variable delays in **Table 4**). Interestingly, it was also common for trained RNNs to fail to generalize despite responding correctly on all training trials (examples in **Fig. S3a**), whereas feedforward models trained on the traditional task format never failed to generalize (examples of outputs in **Fig. S2b**; LR: 100 out of 100 instances; MLP: 100 out of 100 instances; see also [104,105] for additional results in MLPs), highlighting an essential difference posed by the requirement for WM.

**Figure 2:**
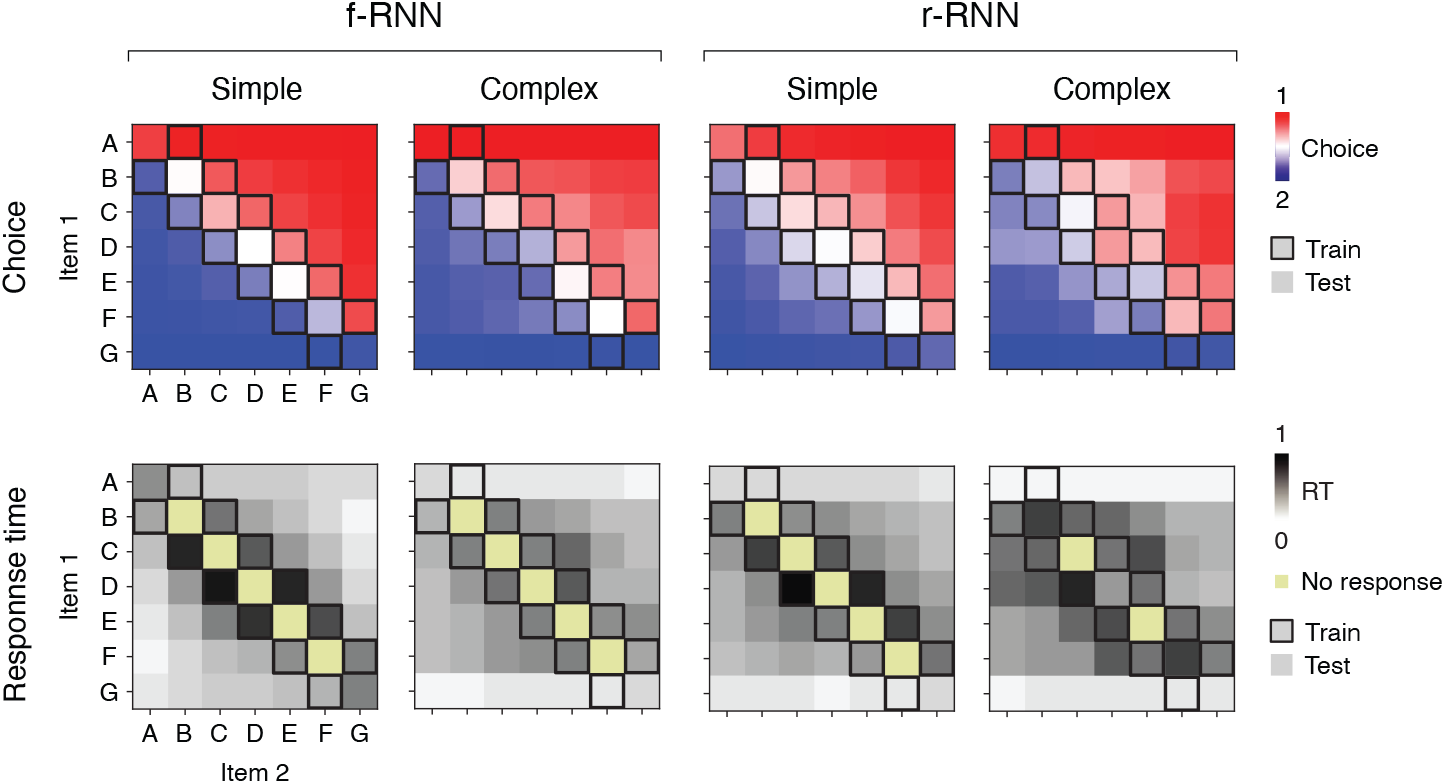
A collection of recurrent neural networks (RNNs) perform transitive inference (TI). Four example RNNs that perform delay TI. Each example RNN is plotted in a column. At left are fully-trainable RNNs (f-RNNs); at right are recurrent-trainable RNNs (r-RNNs); for each, one RNN from each of two training regimes (simple and complex; in particular, simple (high) and complex (high)) are plotted. Top row, network output by trial type. Plotted for each trial type (in squares, defined by items 1 and 2; e.g. AB, BA, AC) is the value of output unit 1 minus that of output unit 2 (Choice, red and blue shades, corresponding to choice 1 and 2, respectively; compare to Fig. 1b). Note RNNs were trained (optimized) only on training trial types (boxed squares). Bottom row, response time (RT) by trial type. In each trial, the choice response of the RNN was defined by the identity of the output unit (linear readout; one for each of two choices) that was first to reach a fixed activity threshold (85%); RT was defined as the time taken for the output unit to reach fixed activity threshold (85%) in the choice period (see **Fig. S1a**), measured as a proportion of the full duration of choice period.

**Table 1:**
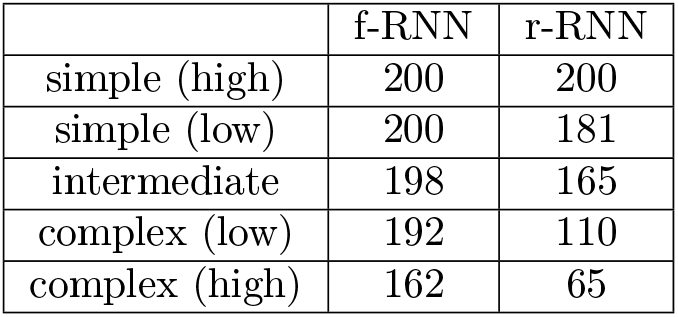
Number of RNNs that fully generalized out of 200 trained instances. All trained instances responded correctly on all training trials, regardless of generalization performance.

Notably, we also observed that, among RNNs, fully-trainable RNNs (f-RNN) and simple training regimes more frequently yielded RNNs that generalized (**Tables 2 and 5**), providing an initial hint that RNN variants might have functionally important differences.

**Table 2:** PCA % variance explained by top 3 PCs across RNNs. Mean ± s.d. (n=65-200 instances / variant; see Table 1).

|  | f-RNN | r-RNN |
| --- | --- | --- |
| simple (high) | $98.1 \pm 0.4$ | $87.2 \pm 1.9$ |
| simple (low) | $97.1 \pm 1.4$ | $88.6 \pm 2.2$ |
| intermediate | $91.5 \pm 1.5$ | $85.5 \pm 2.6$ |
| complex (low) | $82.5 \pm 3.4$ | $79.0 \pm 4.2$ |
| complex (high) | $86.4 \pm 2.8$ | $91.3 \pm 1.7$ |

## 5 RNNs performing TI show multiple emergent behaviors

Decades of work have established that subjects performing TI widely show striking patterns of behavior, manifesting both in performance and response times (RT) [53, 60]. These behaviors are based on trial type, with each trial type defined by items (e.g. AB, BC, AC) and where each item is defined by its position, or “rank”, in the transitive schema (**Fig. 1a**; rank of A is highest, while G is lowest). As recognized previously [60], these empirically observed behaviors are not only important constraints on explanatory accounts of TI, but also potential sources of insight into underlying implementations.

We therefore investigated whether these behaviors (described below), and possibly others, were expressed by RNNs that successfully performed TI (**Fig. 3**). Importantly, expression of these behaviors would effectively be emergent, since the RNNs were neither pre-configured nor trained to express particular behaviors beyond that of responding correctly on training trials. Further, prior work on TI has focused on the traditional TI format (**Fig. 1c**; lacking a WM delay), leaving unclear whether there exist behavioral patterns that explicitly stem from imposing a WM delay.

**Figure 3.**
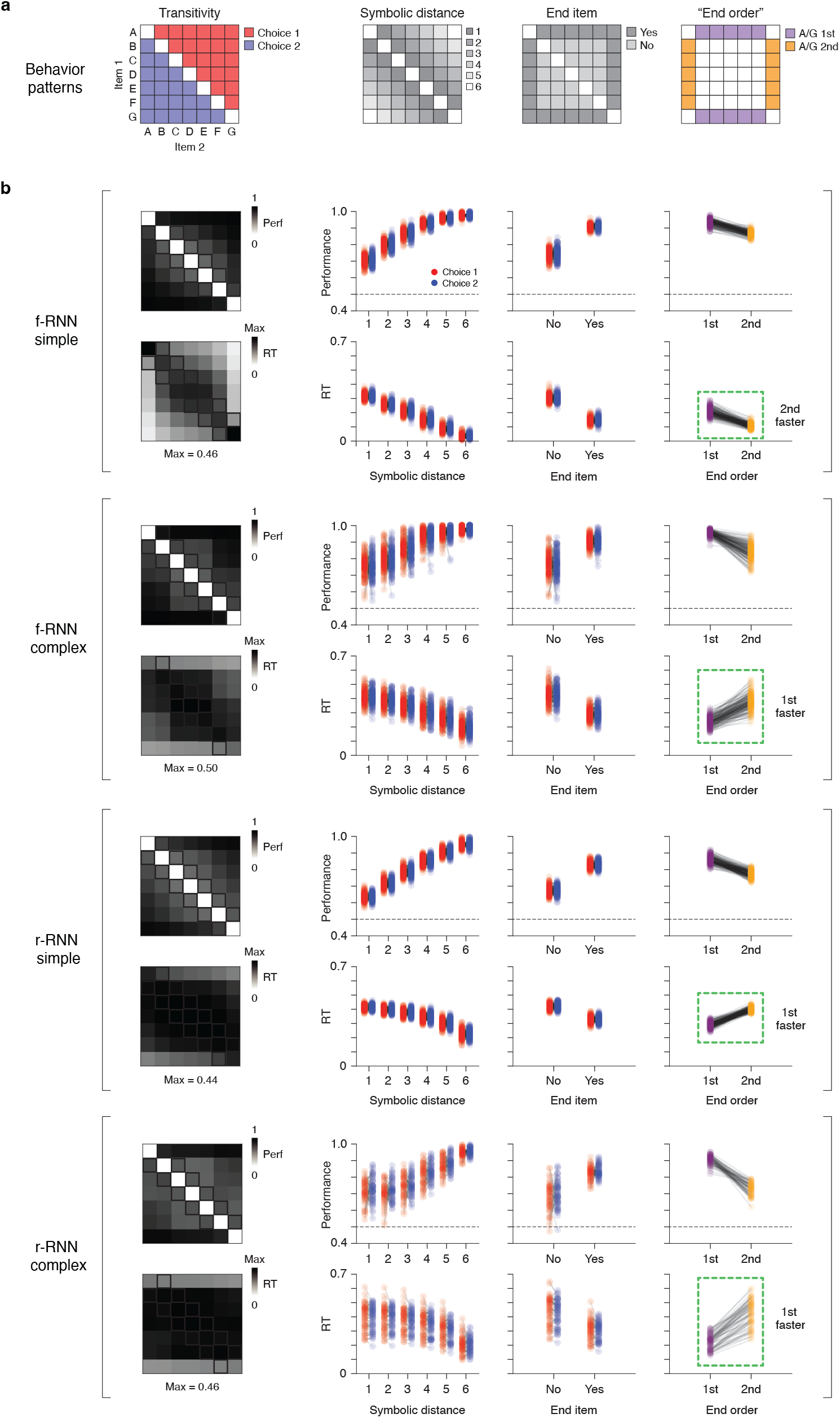
RNNs performing TI show multiple emergent (untrained) behaviors. **a**, Schematic of behavior patterns. Each schematic shows the trial types (items 1 and 2; squares) defining the behavioral pattern. **Transitivity**: the correct choice (item 1 or 2) for transitive inference (i.e. the item ‘higher’ on a transitive schema, Fig. 1a). **Symbolic distance**: the size of the difference in rank between item 1 and 2; rank is an item’s discrete position in the transitive schema (A: 1, B: 2, C: 3, etc.). **End item**: whether the trial type contains an item at either extreme of the transitive schema (‘end item’; here A or G). **End order**: whether the end item (A or G) is the 1st item (item 1) or 2nd item (item 2) presented in the trial; defined only for delay TI (see Fig 1c and **Fig. S1a**). **b**, Behavioral results across four RNN variants (n = 65-200 instances / variant; see **Table 1** for counts). The four RNN variants were defined by different learnable connectivity (f-RNN vs. r-RNN) and training regime (simple vs. complex, corresponding here to simple (high) and complex (high), respectively). Results from each RNN variant are presented in a block of two rows (top row: average performance (proportion correct); bottom row: average RT (proportion of the duration of the choice period; in trial-type matrix (leftmost plot), values are normalized to the maximum value observed across trial types (Max, value reported at bottom)). Column 1: Averages across RNN instances by trial type. Columns 2-4: Averages across trials by trial type, for all RNN instances (500 simulated trials / trial type; each point corresponds an RNN instance). Trial types follow those in panel a (column 2: symbolic distance; column 3: end item; column 4: “end order”) and distinguish between choice 1 vs. choice 2 trial types (red vs. blue, respectively; diagrammed in panel a). Two versions of the “end order” pattern are highlighted (1st- vs. 2nd-faster; dotted green boxes).

Given these aims, we took the following approach to characterize task behavior across RNNs. First, we focused on RNNs that generalized fully (correct responses for every test trial type). Next, for each individual RNN, we simulated the networks with progressively increasing levels of intrinsic noise until the average performance of the RNN was >50% on training trials and <96% on testing trials (i.e. sub-asymptotic performance, the level of performance for which behavioral patterns have been observed). Lastly, we ran simulations (500 runs, all trial types) at this noise level, from which we then measured performance (% correct) and RTs. RTs were measured using a standard criterion (time to a fixed threshold in output units [106]).

We found that RNNs exhibited not only previously established behavioral patterns, but also a novel behavioral pattern not previously studied (**Fig. 3**; additional RNN variants in **Fig. S3c**; analogous analysis in feedforward models in **Fig. S2c-d**). Importantly, behavior was qualitatively and quantitatively comparable to that of living subjects (**Fig. S4**). We address each behavioral pattern in turn.

**Figure 4:**
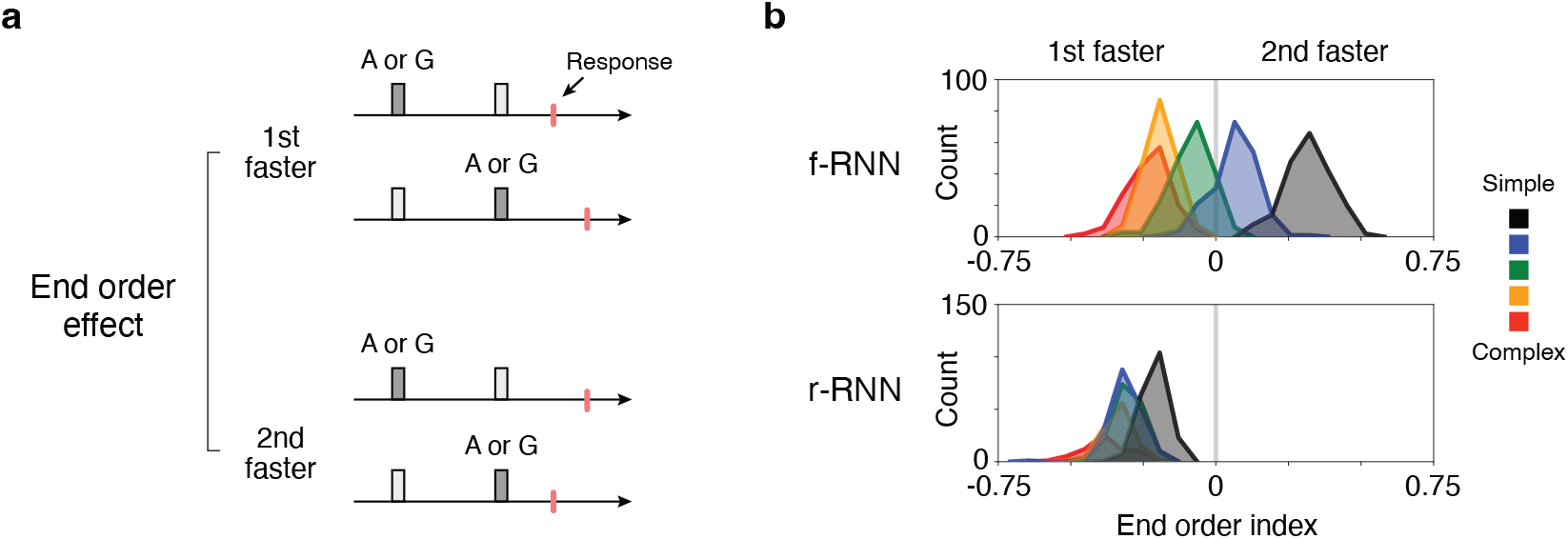
RNNs show a novel order-dependent behavior having two distinct versions. **a**, Schematic of the end order effect, a response-time (RT) behavioral pattern. The behavioral pattern is observed in RNNs in two qualitatively different versions: 1st-faster vs. 2nd-faster. A schematic showing the trial types used to quantify the effect is in Fig. 3a (fourth column). **b**, Histograms of end-order behavior across RNNs (counts: RNN instances). The behavior was quantified as the difference of RTs divided by their sum (end order index; RTs calculated for trials where end items (A and G) occurred either 1st vs. 2nd). RNN variants (f-RNN vs. r-RNN and simple vs. complex training regimes) follow that diagrammed in Fig. 1d. These results summarize those in the last column of Fig. 3.

### The symbolic distance effect

A standard observation across behavioral studies of TI is the “symbolic distance” effect: the larger the difference in rank between items (e.g. AD vs. AB), the higher performance and lower RT [51, 53, 59, 60]. We found that RNNs performing TI invariably exhibited the symbolic distance effect (**Fig. 3**, second column; see also **Fig. S4**).

### The end item effect

Along with the symbolic distance effect, a standard behavioral observation in subjects is the “end item” (or “terminal item”) effect: trials containing either the highest- or lowest-rank item (“end items”; here A and G) are associated with higher performance and lower RTs [53, 60, 107]. We found that RNNs performing TI invariably exhibited the end item effect (**Fig. 3**, third column; see also **Fig. S4**).

### The “end order” effect: a novel behavior with two distinct versions

Traditional TI presents items without an explicit intervening delay, prompting subjects to choose on the basis of item position (e.g. left vs. right designated as item 1 vs. 2, respectively; **Fig. 1b**). In contrast, delay TI prompts subjects to choose on the basis of item order, i.e. whether an item is presented 1st vs. 2nd (before vs. after delay, designated item 1 vs. 2, respectively). Critically, in a subject performing delay TI, this difference in task paradigm may incur an order-dependence in how items are evaluated, i.e. XY and YX trials may differ, neurally and/or behaviorally (besides their different correspondent correct responses). Any such order-dependent effect would be observable in a matrix of trial types (i.e. items 1 and 2) as an asymmetry across the main diagonal.

Unexpectedly, we observed an order-dependent effect that was widespread and also strikingly varied across RNNs: in trials containing an end item (A or G), response times (RTs) were often highly order-dependent (e.g. RTs were faster in AX trials than XA trials). This effect was apparent as an asymmetry in trial-type matrices of RTs (‘stripes’ in the first/last column or first/last row), both in individual RNNs (e.g. **Fig. 2**, bottom row) and in averages across RNNs for a given variant (**Fig. 3b**, first column).

Strikingly, the pattern occurred in two qualitatively different versions. Some RNNs showed lower RTs if an end item (A or G) was presented first rather than second (**Fig. 3**, fourth column, f-RNN simple; also **Fig. 2**, first example), while other RNNs showed the opposite behavioral pattern, namely, lower RTs if an end item (A or G) was presented 2nd (rather than 1st) (**Fig. 3**, fourth column, f-RNN complex, and both r-RNN simple and complex; also **Fig. 2**, second through fourth examples).

We termed the basic shared pattern the “end order” effect (i.e. end-item order; quantified separately in the last column of **Fig. 3**), and its former and latter manifestations as ‘1st-faster’ and ‘2nd-faster’ versions, respectively. In examining this behavior across networks using a quantitative index (end order index, ranging from −1 (1st-faster) to 1 (2nd-faster); see Methods) (**Fig. 4**, additional RNN variants in **Fig. S3d**), we found that the two versions of the behavior systematically differed with respect to both types of RNN variants, whether learnable connectivity (f-RNN vs. r-RNN) or training regime (simple to complex). First, we found that r-RNNs virtually always showed the 2nd-faster pattern (**Fig. 4b**, bottom row), whereas f-RNNs showed both (**Fig. 4b**, top row). Second, we found that complex-regime RNNs nearly always showed the 2nd-faster pattern (**Fig. 4b**, red histograms in top and bottom rows). In contrast, simple-regime f-RNNs consistently showed the 1st-faster pattern (**Fig. 4b**, both simple (high) and simple (low), black and blue histograms, respectively, in top row).

The finding that RNNs performing TI show qualitatively different patterns of behavior (**Fig. 3**, fourth column, **Fig. 4**) furthermore implied that the networks had different underlying neural solutions, motivating direct investigation.

## 6 A simple neural solution to delay TI

Neural recordings in living subjects performing TI indicate that single neurons in the brain can encode variables relevant to TI, including symbolic distance [62, 63]. It remains unclear what collective neural process or activity pattern implements the comparison operation – akin to a ‘>’ operator – that generalizes transitivity to novel combinations of items.

For initial insight, we analyzed how purely feedforward models generalized transitively when WM was not required (logistic regressions and multi-layer perceptrons trained on the traditional TI task format, **Fig. S2**). Examination of unit activations in these models indicated that transitive comparison was implemented using a “subtractive” solution, where the rank (A, B, C, etc.) and position (left vs. right) of each item was mapped to the magnitude and sign, respectively, of unit activation (**Fig. S2e**). Notably, in either feedforward model (LR or MLP), this subtractive solution was sufficiently realized by the (trained) feedforward weights operating directly on the input [104], clarifying that a direct operation on otherwise arbitrary inputs (items A, B, C, etc. encoded as high-dimensional (100-D) random vectors in input activity space) is sufficient to perform TI when feedforward input can be learned or trained. This further raised the question of how TI is performed when modifiable feedforward connectivity is not sufficient (e.g. when WM is required) or not available (e.g. as in r-RNNs).

We thus sought to clarify neural implementations in RNNs performing TI. Given the above findings in feedforward models, we broadly hypothesized that other neural systems having trainable feedforward connectivity — such as f-RNNs — might adopt a similar solution. Beyond this hypothesis, we sought to address three unknowns. First, it was not clear whether or how the subtractive solution observed in feedforward models – which relies on simultaneous input and thus requires twice the number of input connections as RNNs – was relevant to delay TI. Second, it was also not clear whether f-RNNs would consistently adopt the same solution, across either different instances (different random initializations of network weights) or different training regimes (simple vs. complex). Lastly, it remained unclear whether the implementation of TI in r-RNNs, which have no trainable feedforward connectivity, would show any similarity to the implementation of TI in models with trainable feedforward connectivity.

To address these points, we began by investigating the implementation of TI in fully-trainable RNNs (f-RNNs) trained in the simple regime (**Fig. 5**), as we thought these networks might show the most similarity to the purely feedforward models. We first observed that population activity was consistently low-dimensional (% variance explained by top 3 PCs, **Table 2**). Next, in examining population trajectories, we observed two signature patterns (**Fig. 5a**): (1) a **linearly-arranged rank-ordered** response to item presentation (**Fig. 5a**, green-shaded circles, corresponding to the network response to item 1), and (2) a **rotation** during the delay period (**Fig. 5a**, bottom), suggesting an oscillatory dynamic.

**Figure 5:**
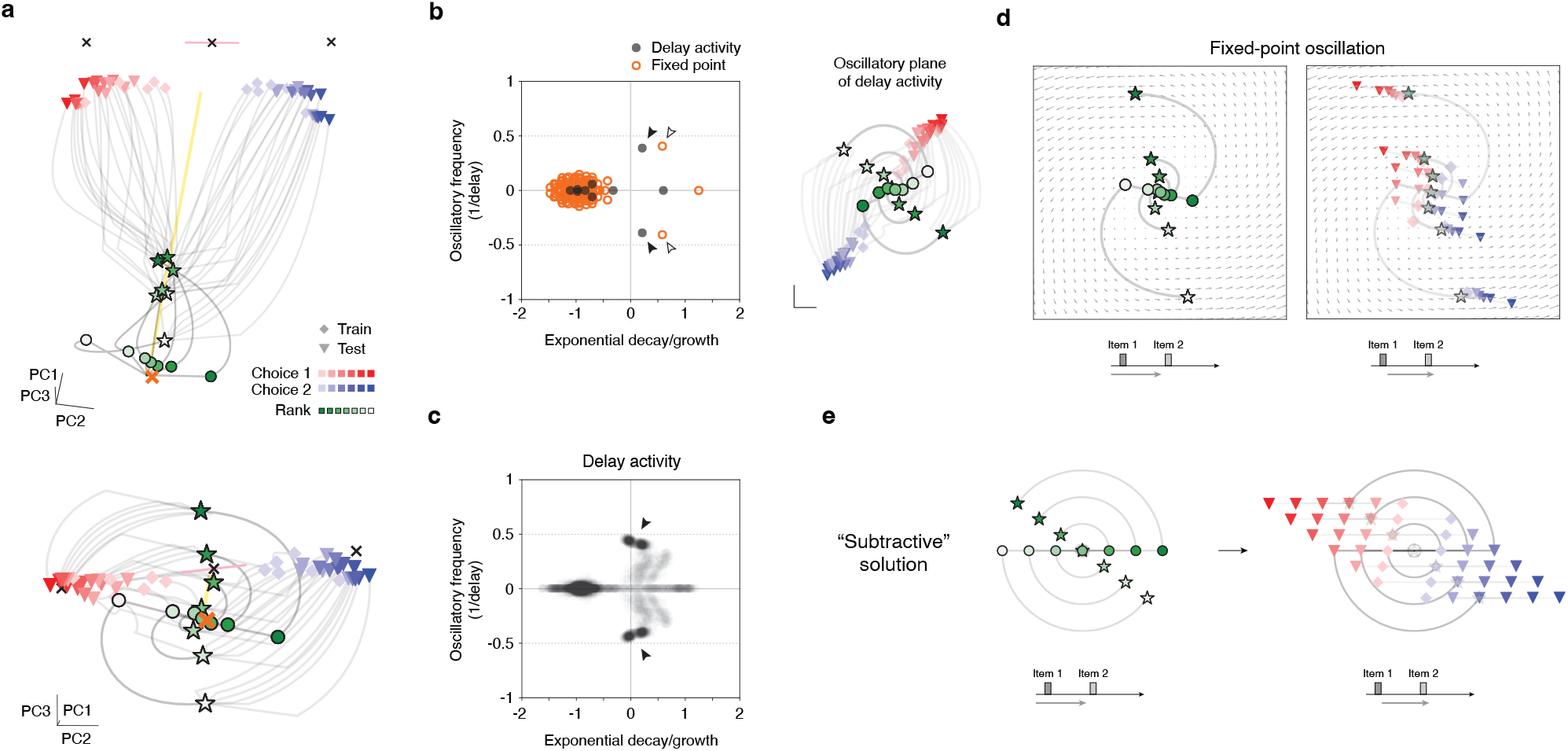
A simple neural solution to delay TI. **a**, Population activity trajectories in an example RNN (simple-regime f-RNN) that performs TI. Top and bottom plots show two different views. Shown are trajectories from all 42 trial types (see Fig. 1b). To clarify the operation of the network, three trial times are highlighted: (i) presentation of item 1 (green circles; shade indicating item rank: A (dark green) to G (white)), (ii) the last time point of the delay period (green stars; same color convention), (iii) last time point of the trial (red/blue symbols; red: choice 1 trials, blue: choice 2 trials, light to dark shading indicating symbolic distance (1 to 6); diamonds: training trials, triangles: test trials). Also shown: cross-condition mean (XCM; the average trajectory across all trial types) (yellow line) and fixed points (FPs) (crosses). Two FPs were attractors (lone black crosses), one FP was a saddle point (black cross with pink line indicating axis of unstable mode), and one FP (orange cross) was located near trajectories during the delay period (‘early-trial’ FP). Note two prominent activity patterns: linearly arranged rank-ordered activity upon presentation of item 1 (green circles) and the oscillatory evolution of trajectories in the delay period (circles to stars). **b**, Linear dynamics of RNN in panel a. Left, eigenvalue spectra of the RNN. The spectra were calculated in two ways: first, from delay-period neural activity (black points; inferred via least-squares linear fit, *R*^2^ = 0.65) and second, from linearization of the network with respect to the early-trial FP (orange circles; FP shown as orange cross in panel a). Right, population activity trajectories of the RNN plotted in the plane of the oscillation inferred from delay period activity (filled arrowheads in spectra plot). **c**, Linear dynamics of simple-regime f-RNNs (n=400 instances; 200 simple (high) and 200 simple (low)). Eigenvalue spectra of delay-period neural activity (grey translucent points; inferred via least-squares linear fit, *R*^2^ ~0.6-0.9; see **Fig. S5a**). Note the density of oscillatory modes with frequency ~0.5 cycles / delay (filled arrowhead). **d**, Activity trajectories in the oscillatory mode of the linearized RNN. The oscillatory mode is that of the linearization of the early-trial FP (open arrowheads in panel b). Plotted in background are flow field vectors (not to scale; shown for motion direction). To clarify how the activity evolves, trajectories are plotted in two stages of the task trial (left and right panels; schematic at bottom of each): early trial (left) and presentation of item 2 (right). Three trial times are highlighted: (i) presentation of item 1 (green circles; shade indicating item rank: A (dark green) to G (white)), (ii) the last time point of the delay period (green stars; same color convention), (iii) presentation of item 2 (red/blue symbols; red: choice 1 trials, blue: choice 2 trials; diamonds: training trials, triangles: test trials). Note the separation of choice 1 vs. 2 trials (red vs. blue symbols), indicating that correct responses were evolved in the activity space. **e**, Diagram of solution expressed in networks: population-level “subtraction”. Plotted are activity trajectories generated by simulating a 2D linear dynamical system defined by an oscillation of frequency ~0.5 cycles / delay, with initial condition at the origin and input vectors encoding task items (A, B, C, etc.) in ordered collinear arrangement in state space (compare to panels a and d). Trial-based input (item 1 - delay - item 2, see Fig. S1a) was applied to the system. Plotting conventions are the same as in panel d. For greater detail of the solution, see **Supplementary Diagram**.

Pattern 1 suggested a similarity to the subtractive solution, as the pattern was characterized by an intrinsically 1D activity structure across trial types (observed in single units in feedforward models, whereas population activity space in RNNs) and was due to trained feedforward connectivity. Pattern 2 suggested that a low-dimensional dynamical process — potentially a single (2D) oscillation — might be sufficient to account for how these networks implemented transitive comparison across time. Indeed, we found that RNN activity during the delay period was effectively described by linear dynamics (ordinary least-squares fit) characterized by an oscillatory mode correspondent with pattern 2 (**Fig. 5b-d**; *R*^2^ ~0.6-0.9 in simple-regime (high) and simple-regime (low) f-RNNs, **Fig. S5a**). Given these clues, we then sought to identify the underlying solution explicitly. Importantly, prior work has found that analysis of population-level neural dynamics in trained RNNs with respect to fixed points (FPs) can identify dynamical components that have specific functional roles [84,108–110] and are jointly sufficient to perform cognitive tasks.

Taking this approach, we found that simple-regime f-RNNs had an “early-trial” fixed point located near activity trajectories at the beginning of trials (**Fig. 5a**, orange cross). This fixed point had an oscillatory mode of frequency ~0.5 cycles/delay (highlighted by open arrowheads in **Fig. 5b**) that was orthogonal to the average trajectory across trial types (i.e. the cross-condition mean (XCM), a population activity component consistently observed across trial-based tasks [111–113]; plotted as yellow line in **Fig. 5a**; analysis of orthogonality in **Fig. S8b-c**). Some RNN instances also showed additional FPs that were overtly associated with other task functions (**Fig. 5a**, black crosses): two “choice” FPs that are stable FPs (attractors) toward which trajectories corresponding to each of the two choices travelled and would eventually settle following item 2 presentation (not shown), and a saddle FP, located between the two choice FPs, that was stable except for a single unstable axis oriented toward each choice FP (**Fig. 5a**, choice FPs: black crosses respectively near end of choice 1 and choice 2 trajectories; saddle FP: center black cross, saddle unstable axis in pink line). This unstable axis suggests a “choice axis,” as it was aligned with the segregation of activity trajectories with either of the two choice FPs following presentation of item 2 (**Fig. 5a**, item 2 presented at the time point corresponding to the shaded stars), and was parallel to the direction between choice 1 and choice 2 activity states. This latter pattern of activity indicates a choice axis (direction pointing from choice 1 vs. choice 2 activity states; see Methods) that is parallel to the oscillatory mode (analysis of angle in **Fig. S8b-c**). Together, these dynamical components indicated that the transitive comparison could be performed within the 2D subspace (plane) of the oscillatory mode of the early-trial FP. To test this possibility, we evaluated whether TI could be performed solely by activity and dynamics within the subspace of the identified oscillation (constituting a linear approximation of the oscillation seen in the RNN; see Methods for identification procedure). We found that this oscillation, when presented with trial-structured input (item 1 - delay - item 2), yielded activity trajectories for which correct choice was linearly separable (**Fig. 5d**).

This clarified the implementation of transitive comparison over time in these networks (**Fig. 5e**): namely, a single oscillation that uniformly re-orients activity states corresponding to item 1 (A, B, C, etc.), doing so via a common angular displacement. In the delay period, following the presentation of item 1, this re-orientation serves to shift the activity state in the direction opposite to the direction of the activity displacement due to item presentation. Note that the activity displacement due to item presentation is rectilinear – a consequence of its implementation as feedforward input – whereas the dynamics-driven activity displacement in the delay period is angular. The re-oriented activity state is thereby “subtractive” with respect to the activity shift subsequently elicited by the presentation of item 2.

Notably, a particular angular shift of ~0.5 cycles (resulting from an oscillatory frequency of ~0.5 cycles/delay) re-orients the activity state to be opposite (diametric) to that imposed by the presentation of the item prior to the delay. This can be likened to a change-of-sign along a “comparison” axis in population activity space (**Fig. 5e**), and is analogous to the mapping of signs (− vs. +) to item position (left vs. right) in the subtractive solution to TI in the feedforward models (**Fig. S2e-f**); this solution can be thought of as subtraction at the population level (**Fig. 5e** and **Supplementary Diagram**). The resulting activity along the comparison axis accounts for transitive comparison and the symbolic distance effect, as long as the output implemented by the network (e.g. the readout weights of the output units) is aligned with the comparison axis.

## 7 Geometric signatures of networks performing delay TI

The above findings identify a simple (population-level) neural solution to TI (**Fig. 5**), but do not address other possible neural solutions. Importantly, we earlier observed that RNN variants performing TI showed qualitatively different behavioral patterns (**Fig. 4**), implying that there exist alternative neural solutions.

To clarify these solutions, we took the subtractive solution as a point of reference. We sought to identify one or more activity patterns that characterize this solution while also being easily quantifiable, whether in neural models or in putative experimental data. Such characteristic or signature activity patterns could then be a means by which to survey and/or discover alternative solutions. From the simple solution (**Fig. 5** and **Supplementary Diagram**), two component activity patterns were (1) the linearly-arranged rank-ordered arrangement of activity states (pattern 1), which here refer to as “ordered collinearity,” and (2) the oscillation associated with transitive comparison (pattern 2). Across RNNs, we assessed these two patterns in turn.

Surprisingly, we found that pattern 2 was not unique to the subtractive solution. A striking indication that this was the case came from analyzing r-RNNs (recurrent-trainable RNNs), an RNN variant that necessarily adopts some form of an alternative solution, since the subtractive solution depends on the ordered collinear activity made possible by learned feedforward connectivity (not available in r-RNNs by definition). Despite this fundamental difference, neural activity in r-RNNs appeared remarkably similar to that of simple-regime f-RNNs (**Fig. S6a**, compare to **Fig. 5a**): in particular, r-RNNs showed an on oscillatory mode consistent with pattern 2 (frequency ~0.5 cycles/delay; identified either from fixed-point linearization or from linear dynamics inferred from delay period activity, **Fig. S6b-c**). Further, as in the case of simple-regime f-RNNs seen earlier (**Fig. 5d**), analysis of the linearized dynamics revealed that this single oscillation, when combined with the task inputs (item 1 - delay - item 2), was sufficient to accomplish transitive comparison (**Fig. S6d**). Interestingly, this sufficiency was only apparent when the oscillation was applied to full-dimensional activity (**Fig. 6d**, bottom row; further explanation in Methods) and not activity reduced to the 2D subspace of the oscillation (**Fig. 6d**, top row), an indication that the implementation of TI in these networks also depends on higher-dimensional activity components. These results indicate that though r-RNNs cannot adopt the subtractive solution (**Fig. 5e**), the implementation in r-RNNs nonetheless can rely on a single dynamical pattern (here an oscillation) to perform delay TI.

**Figure 6:**
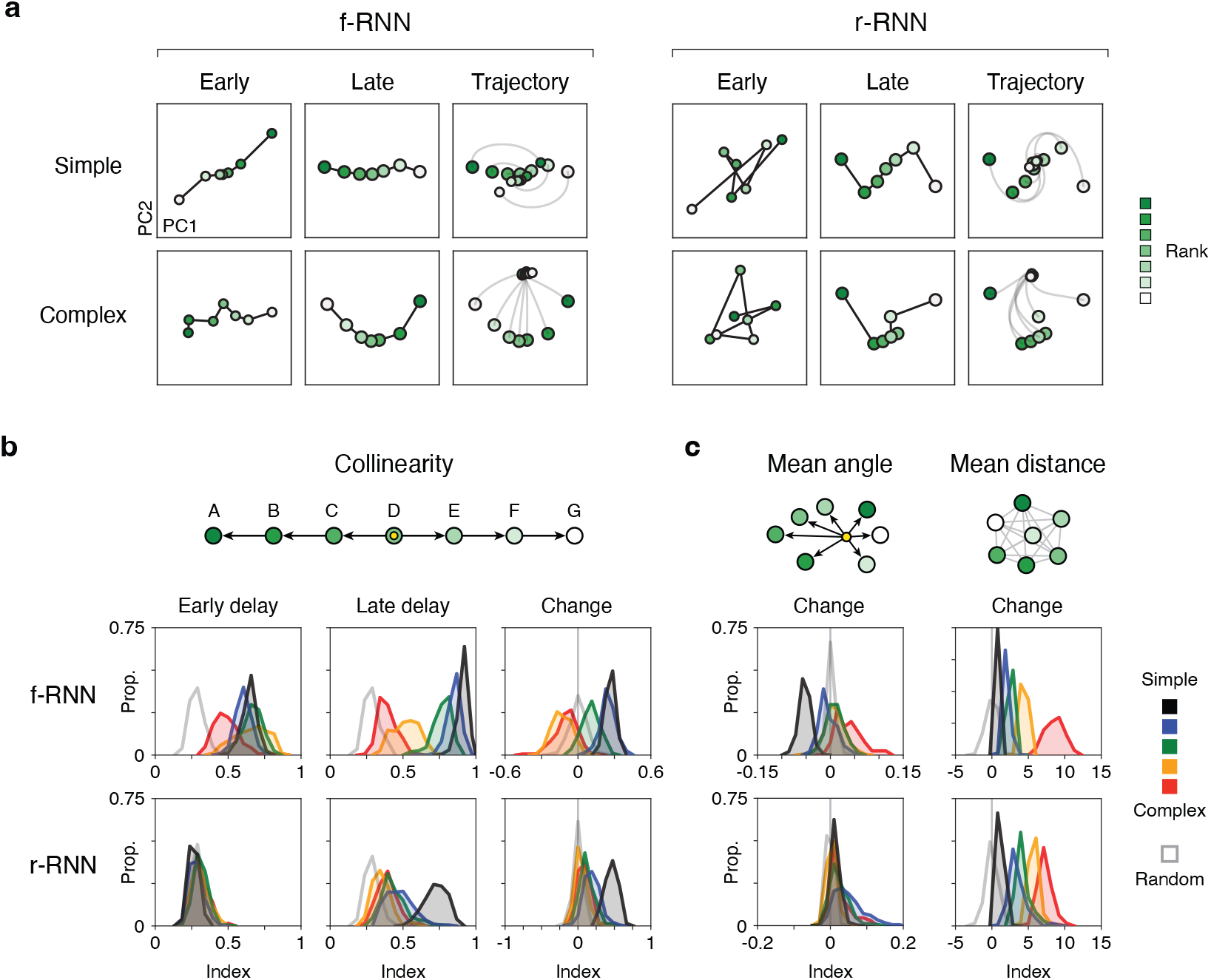
Geometric signatures of networks performing delay TI. **a,** Delay period population activity in four example RNNs. In each example, the seven trial types corresponding to different identities of item 1 are shown (item rank indicated by green shade). PCs were calculated from population activity at the end of the delay. Axes units differ between plots to show the geometric arrangement at each time point. Small vs. large circles indicate early vs. late delay activity, respectively. **b**, Collinearity: schematic (top) and quantification across RNNs performing delay TI (bottom). Top, schematic of geometric arrangement in population activity space. The corresponding index (collinearity index; see Methods) quantifies whether neural activity during the delay matches this geometric arrangement. Angles were measured relative to the cross-condition mean (XCM, yellow circle; shown in example RNNs in Figs. 5a and 6a). Bottom, histograms of index values across RNNs. Index values were measured in three respects (columns): early delay, late delay, and change across delay (late - early). Early and late delay were defined here as the first and last timesteps of the delay, respectively; similar results were obtained when using averages from the first and last quarter of the delay. **c**, Mean angle and mean distance: schematics of each measure (top) and histograms of index values across RNNs (bottom). For both mean angle and mean distance, all pairwise angles and pairwise distances, respectively, were calculated in population activity space (between trial types based on item 1, i.e. A, B, C, etc.; green-shaded circles) and averaged. Angles were measured relative to the cross-condition mean (XCM; yellow circle). For both activity geometries, the change during delay (late - early) is plotted. All plots show histograms of instances for each RNN variant (n=65-200 instances / variant; see **Table 1**), in addition to randomly generated data (open grey histograms); all quantifications were performed in the top 10 PCs of neural activity during the delay (quantification in top 2 PCs presented in **Fig. S7b**).

Unlike pattern 1, pattern 2 – ordered collinearity, a simple geometric pattern in population activity space (**Fig. 5a, c**) – definitively differed across RNNs performing the task. As an initial assessment, we examined activity in activity space during the delay, which reiterated that r-RNN activity was not collinear at the start of the delay (Early plots, r-RNN; **Fig. 6a**, **Fig. S7a**), as expected given the lack of learnable feedforward connectivity in r-RNNs. Notably, RNNs not trained in the simple regime expressed a variety of non-collinear patterns in activity space; in particular, networks often expressed a “V” shape, where the activity corresponding to the items along the transitive schema is “folded” or curved in activity space (f-RNN complex in **Fig. 6a**; additional examples in **Fig. S7a**) similar to that observed in recent studies that involve learning an explicitly-cued transitive relationship between stimuli [105, 114].

To assess collinearity across models, we defined quantitative indices for both collinearity and ordered collinearity (ranging from 0 to 1 for collinearity and −1 to +1 for ordered collinearity, schematic in **Fig. 6b**; ordered collinearity presented in **Fig. S7b**). Each index is measured in the full population activity space of individual networks (ambient N-D space, reduced here to top 10 PCs to aid comparison to predominant responses in neural data).

As expected from previous observations (**Fig. 5** and **S6**), simple-regime f-RNNs expressed collinearity index values that were relatively high (>0.5) or near 1 (**Fig. 6b**, black and blue histograms in upper row, first and second columns; collinearity, early delay: simple (high): 0.65 ± 0.05, simple (low): 0.60 ± 0.05; collinearity, late delay: simple (high): 0.91 ± 0.03, simple (low): 0.85 ± 0.05; mean ± s.d., n=65-200 for each variant, see **Table 1**) and that were virtually always higher than that of r-RNNs at the start of the delay (**Fig. 6b**, first column; r-RNN: 0.29 ± 0.05, n=921 instances) or of randomly generated activity vectors (random vectors: 0.28 ± 0.05, n=200 draws; equivalent to r-RNNs at the start of the delay).

Interestingly, complex-regime f-RNNs expressed significant collinearity at the start of the delay (early delay). In these networks, index values were lower than that simple-regime f-RNNs, yet consistently higher than that of random activity (**Fig. 6b**, red histograms in upper row, first and second columns; collinearity, early delay: complex (high): 0.49 ± 0.09, vs. simple (high) and simple (low), p < 10*^−^*^49^, vs. random p < 10*^−^*^57^; collinearity, late delay: complex (high): 0.39 ± 0.06, vs. simple (high), p < 10*^−^*^76^, vs. random, vs. random p < 10*^−^*^42^; rank-sum tests for comparisons; see **Table 1** for counts). The expression of significant collinearity at the start of the delay indicates that feedforward connectivity in these networks was modified in training.

Lastly, we found that, in parallel to f-RNNs, different training regimes in r-RNNs yielded consistently different activity geometries, thus making it possible to distinguish these RNN variants on the basis of neural activity. In particular, complex- vs. simple-regime r-RNNs expressed overtly different degrees of collinearity at the end of the delay period (>0.5 vs. <0.5 in simple vs. complex, respectively; **Fig. 6b**, black vs. red histograms in bottom row, second column; simple (high): 0.73 ± 0.08, complex (high): 0.39 ± 0.07, p < 10*^−^*^32^).

These several results establish that collinearity, a population-level activity geometry quantifiable for any neural system performing delay TI, can be used to distinguish between neural models; the predictions for each of the models are summarized in **Table 3**, with corresponding implications discussed further below.

**Table 3:** Main predictions. ••, a prediction unique to this model. •, two predictions that jointly are unique to this model. “End order” effect: **Fig. 4**. Collinearity: **Fig. 7a**. Mean angle change: **Fig. 7b**. Oscillation in delay period: **Figs. 5c, 6c, S8**. Choice axis encoding: **Fig. 7c**. All neural predictions refer to the top PCs of population activity.

|  | Behavior | Neural activity |  |  |  | Implication |
| --- | --- | --- | --- | --- | --- | --- |
|  | “End order”<br>effect | Collinearity | Mean<br>angle<br>change | Oscillation<br>in delay period | Choice axis<br>encoding |  |
| f-RNN<br>simple | ●●2nd-faster | ●Early: $>0.5$<br>●Late: $>0.5$<br>Change: $>0$ | ●●$<0$ | $\sim 0.5$ cycles/delay<br>XCM cosine: $\sim 0$<br>Choice cosine: $\sim 0.5$ | ●●2nd-dominant | Feedforward<br>input learned,<br>“subtractive”<br>solution |
| f-RNN<br>complex | 1st-faster | Early: $>\text{random}$<br>Late: $>\text{random}$<br>●●Change: $<0$ | $>0$ | n/a | 1st-dominant | Feedforward<br>input learned |
| r-RNN<br>simple | 1st-faster | ●Early: random<br>●Late: $>0.5$<br>Change: $>0$ | $>0$ | $\sim 0.5$ cycles/delay<br>XCM cosine: $\sim 0$<br>Choice cosine: $\sim 0.5$ | 1st-dominant | Recurrent<br>dynamics learned,<br>single oscillation<br>sufficient |
| r-RNN<br>complex | 1st-faster | ●Early: random<br>●Late: $<0.5$<br>Change: $>0$ | $>0$ | n/a | 1st-dominant | Recurrent<br>dynamics learned |

Measuring collinearity yielded further insight into activity geometry across implementations. In examining collinearity index values at the beginning and end of the delay (early and late delay; **Fig. 6b**, left and middle columns), we noticed that index values could increase dramatically across the delay period (e.g. ~0.3 to ~0.8 for r-RNNs trained in the simple (high) regime, **Fig. 6b**, black histograms in bottom row; also previously observable as an “unfolding” of activity trajectories during the delay period in **Fig. S6a**; examples of evolution of angles in **Fig. S7c**). This observation led us to examine the degree to which collinearity changes across the delay (**Fig. 6b**, right column). Intriguingly, f-RNNs exhibited systematic differences depending on training regime: in particular, simple-regime networks invariably increased collinearity during the delay (**Fig. 6b**, black and blue histograms in top row; collinearity change: simple (high): 0.27 ± 0.04, simple (low): 0.25 ± 0.06; mean ± s.d., see **Table 1** for counts), whereas complex-regime networks consistently decreased collinearity during the delay (**Fig. 6b**, both complex (high) and complex (low), red and orange histograms, respectively, in top row; collinearity change: complex (high): −0.10 ± 0.10, complex (low): −0.13 ± 0.08; mean ± s.d.). These qualitatively different patterns of activity (increasing vs. decreasing collinearity) were consistent with differing underlying solutions of TI in these network variants, as implied previously by the distinct patterns of behavior expressed by these variants (**Fig. 4**). More broadly, these findings suggested that, apart from collinearity, there might exist other geometrically defined activity patterns, expressed during the delay period, distinguishing alternative implementations of TI.

Leveraging this insight, we identified one such geometric activity pattern. We were struck by the fact that alternative implementations of TI – namely, the subtractive solution (**Fig. 5f**) vs. the solution in simple-regime r-RNNs (**Fig. S6**; lacking ordered collinearity from feedforward input) – nonetheless both depended upon a similar dynamical pattern: a single oscillation enabling transitive comparison over time. We conjectured that a single oscillation could be a relatively simple case in a broader set of dynamical patterns enabling transitive comparison over time, and, further, that the essential operation of this wider set of dynamics is rotation.

Given this possibility, we hypothesized that activity geometry based on angular rather than distance relationships (i.e. angles vs. distances between activity states in A vs. B trials, A vs. C, B vs. C, etc.) might distinguish between alternative implementations of TI. To test this possibility, we evaluated two geometric measures – mean angle and mean distance (schematic in **Fig. 6c**; see Methods) – that were more general than collinearity indices in characterizing geometric relationships of neural activity. Strikingly, we found that change in mean angle (**Fig. 6c**, left column), but not in mean distance (**Fig. 6b**, right column), categorized RNN variants (values <0 vs. >0) in overall accordance with the two types of alternative implementations previously implied: (1) the subtractive implementation exhibited in simple-regime f-RNNs vs. the implementation in r-RNNs (**Fig. 6c**, left column, black histogram in top row vs. histograms in bottom row; simple (high) f-RNN: −0.052 ± 0.012 (mean ± s.d., vs. 0, p < 10*^−^*^34^; r-RNN, all variants: 0.019 ± 0.029, n=921 instances, vs. 0, p < 10*^−^*^81^; signed-rank tests), and (2) the alternative implementations of simple- vs. complex-regime f-RNNs (**Fig. 6c**, left column and top row, values <0 vs. >0 in black and blue vs. orange and red histograms; simple-regime f-RNNs: −0.052 ± 0.012, n=200 instances, vs. 0, p < 10*^−^*^34^; complex-regime f-RNNs: 0.036 ± 0.027, n=162 instances, vs. 0, p < 10*^−^*^27^; signed-rank tests). Direct assessment of activity geometry in relation to behavior (**Fig. S8a-b**) indicated that mean angle change had a predictive relationship to behavior which was invariant across all RNNs – in particular, positive and negative mean angle changes predicted 1st- vs. 2nd-faster end order behavior, respectively (**Fig. S8a-b**).

We lastly sought to identify specific activity geometries yielding a positive mean angle change. Notably, the “V” activity geometry, which we observed to occur in complex-regime RNNs (in the top 2 PCs, **Fig. 7a**, **Fig. S7a**) and which has been observed in neural responses in related tasks [105, 114], was expressed in networks having a positive mean angle change (see **Fig. S7a**). We further found that this correspondence was observable within the top 2 PCs (**Fig. S7c**). The “V” pattern thus constitutes an activity geometry that is elicited by a positive mean angle change, is associated with 1st-faster behavior, and, moreover, is consistent with empirical observations in related task paradigms.

**Figure 7:**
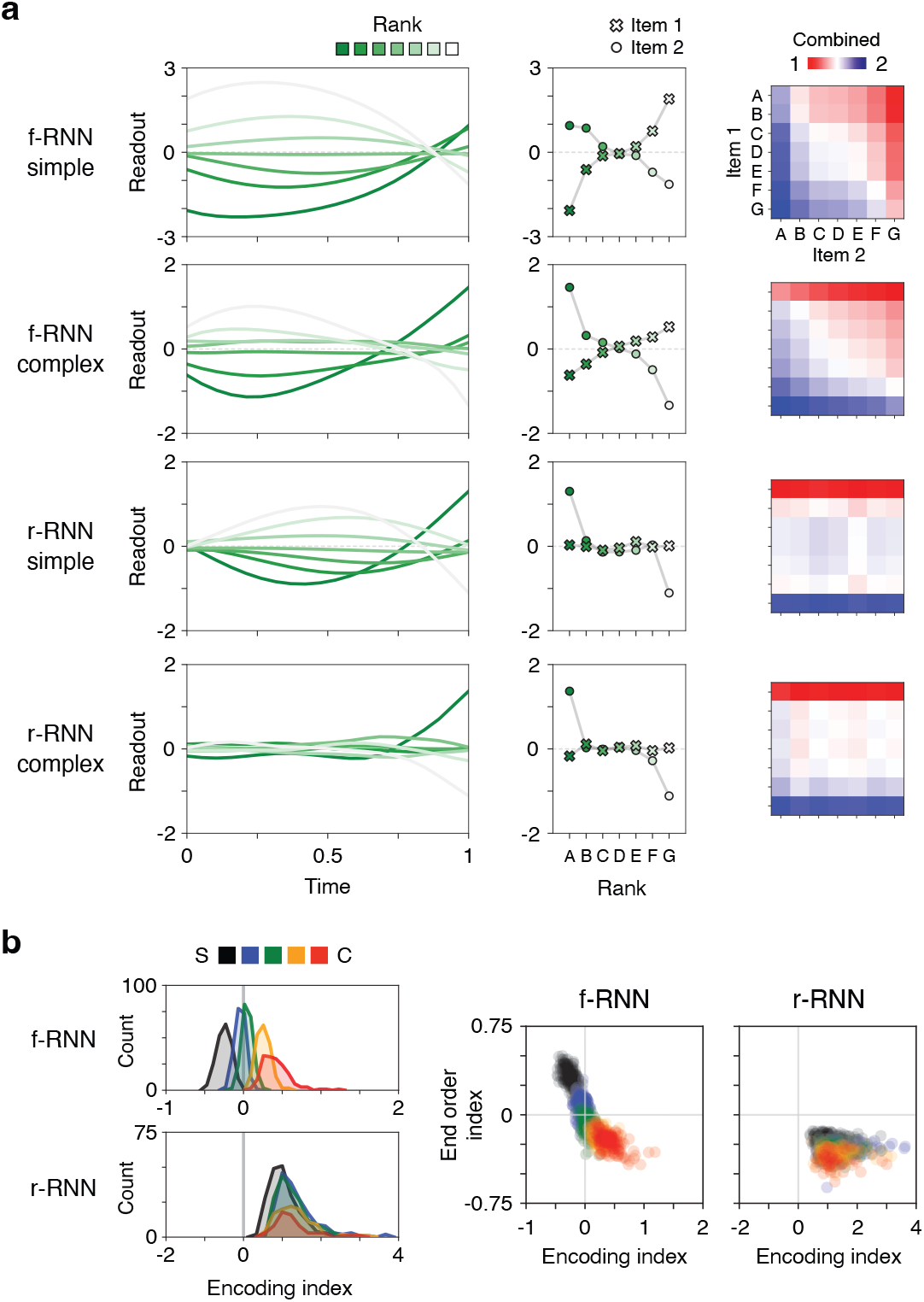
A neural basis for order-dependent behavior. **a,** Neural encodings (axis-projected activity) in four example RNNs (rows). Left column, projection of neural activity along the readout axis (output unit weights). Middle column, item 1 vs. 2 encoding, corresponding to projected neural activity at the end and beginning of the delay, respectively. Right column, sum of item 1 and item 2 encodings. Note that the sum yields large values for trials containing end items, either when the end item is item 1 (rows; examples 2-4) or item 2 (columns; example 1). **b**, Left, histograms of the encoding index across RNNs. The encoding index was defined as the multiplicative gain in the magnitude of end items’ neural encodings (>0: item 1 encoding larger than item 2, ‘1st-dominant’; <0: item 2 encoding larger than item 1, ‘2nd-dominant’). Right, end order index vs. encoding index across RNNs. Note that 1st- vs. 2nd-dominant encodings correspond to 1st- vs. 2nd-faster behavior (end order index <0 vs. >0, respectively). The analogous analyses for the choice axis are presented in **Fig. S8c**.

## 8 A neural basis for order-dependent behavior

The observation that RNN variants performing delay TI express uniquely different activity geometry (**Fig. 6**) corresponding to qualitatively different behavior – i.e. the end order behavior (**Figs. S8a-b**) – suggests that this behavior may have a readily identifiable neural basis.

To investigate this possible neural basis, we conjectured that the two distinct versions of the end order behavior, i.e. the 1st-faster vs. 2nd-faster versions (**Fig. 4**), could be due to a simple difference in how networks internally represent (encode) item 1 vs. item 2. We examined encodings along two axes in activity space expressly relevant to behavior: (i) the readout axis (linear readout weights of output units), the axis along which activity directly generates a behavioral response, and (ii) the choice axis (the direction in activity space from choice 1 to choice 2 trials, measured at the beginning of the choice period; see Methods), a behaviorally relevant axis that can be estimated from neural activity. In initial analyses, we found that the choice axis effectively approximates the readout axis in all RNNs that performed delay TI (**Fig. S5c**).

Examining neural activity projected along either axis revealed two different patterns of encoding across RNNs (**Fig. 7a**), with each pattern varying systematically across RNN variants (**Fig. 7b**). Projections of activity on the readout axis are shown in example RNNs in **Figure 7a**. In a subset of RNNs, we found that when item 1 is an end item (A or G), it is re-encoded during the delay period such that its encoding magnitude is reduced (**Fig. 9a**, upper row). Conversely, in complex-regime f-RNNs and r-RNNs, recoding in the delay period instead increased the encoding magnitude of end items (**Fig. 9a**, second to fourth rows). These two encoding patterns imply that, for trials containing an end item, either item 1 or item 2 is dominant in generating behavior. The difference between the the former (item 1 dominant) versus latter (item 2 dominant) was illustrated by directly summing the encoding values of item 1 and 2 (**Fig. 9a**, right panels). We refer to these two encoding patterns as “1st-dominant” and “2nd-dominant,” respectively. Importantly, these encoding patterns can be quantified in any neural system performing delay TI by evaluating the choice-axis projection of neural activity taken from the end vs. beginning of the delay period, which provide estimates of the encoding value of item 1 (due to recurrent input) and item 2 (due to feedforward input), respectively. We quantified the relative encoding magnitude of item 1 vs. item 2 as a ratio (encoding ratio; item 2 : item 1), doing so for both the readout axis (presented in **Fig. 7**) and choice axis (presented in **Fig. S8c**, yielding qualitatively similar results),

Quantification of the encoding ratio across RNNs revealed a systematic difference predictive of end-order behavior. Across f-RNNs, simple- vs. complex-regime networks showed 2nd- vs. 1st-dominant encodings, respectively (**Fig. 9b**, upper histograms). Strikingly, r-RNNs invariably showed the 1st-dominant encoding (**Fig. 9b**, bottom histogram). Across all networks, these encodings effectively predicted end order behavior (**Fig. 9c**). Given these findings, the two qualitatively different encoding schemes (1st vs. 2nd-dominant) suggest a plausible and experimentally testable neural basis of the end order behavioral pattern (summarized as a neural prediction in **Table 3**).

It is also worth highlighting that the 1st vs. 2nd-dominant encoding patterns correspond to the relative magnitudes of recurrent vs. feedforward input; in this regard, simple-regime f-RNNs were exceptional among RNN variants in expressing stronger feedforward-based encoding. As complex-regime f-RNNs showed stronger recurrent-based encoding, training regime (simple vs. complex) thus determines the respective contributions of feedforward vs. recurrent input to a behaviorally relevant representation in these neural systems.

## 9 Delay TI in human subjects

The above investigation establishes the existence of qualitatively different neural solutions to TI when WM is required, a finding that stemmed from imposing an intervening delay between item presentations (**Fig. 1c**, **Fig. S1**). Yet despite the extensive literature on TI [53, 58], prior studies generally do not impose such a delay (though see [81] for a probabilistic version of TI): rather, items are presented either simultaneously (no delay, **Fig. 1c**; e.g. left and right images on a screen) or as encountered at the discretion of the subject (e.g. odors in separate containers, conspecifics in separate chambers [52, 67, 68]). Consequently, there is a lack of experimental data for testing predictions pertaining to the WM delay, particularly predictions that distinguish between models (e.g. the end order pattern, **Fig. 4**); more broadly, experimental data is also needed to evaluate whether standard TI behaviors are expressed when WM is explicitly required.

We therefore tested delay TI in humans in a large-scale experimental study (392 subjects) using Amazon Mechanical Turk, an online platform enabling testing human subjects on cognitive tasks. As with the neural models, a panel of 7 arbitrary items (fractal images, corresponding to items A to G; **Fig. 8a**) was used; further, within each trial, an intervening delay period (1 sec) was imposed between item presentations (**Fig. 8b**). In each trial, subjects chose either item 1 or 2 via key press (“D” or “K” key, corresponding either to item 1 or 2), and were informed of trial outcome (correct/incorrect) immediately upon choosing. To promote task engagement, subjects were incentivized to select correctly so they could earn performance-based bonus money. Training was performed for a fixed number of trials (training phase; 144 trials consisting solely of training trial types, i.e. A vs. B, B vs. C, etc., divided into 3 blocks of trials), after which testing was conducted (testing phase; 256 trials consisting of both training and testing trial types, divided into 6 blocks of trials). Upon completion of testing, subjects were asked to describe how they performed the task.

**Figure 8:**
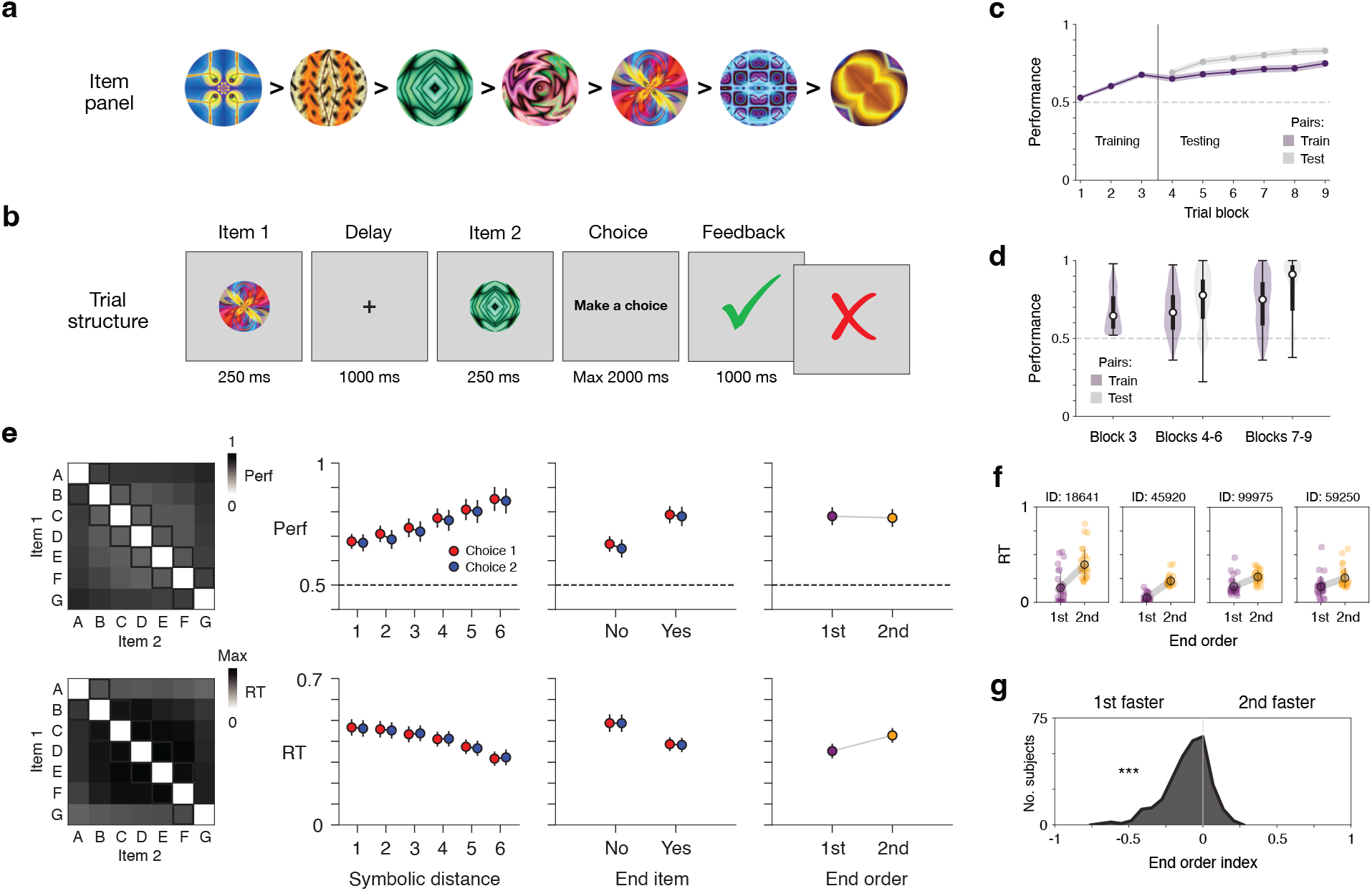
Delay TI in human subjects. **a**, Item panel. Each item is a fractal image. Shown is an example transitive ordering of images; for each subject, the order was randomly generated. **b**, Schematic of trial structure. **c**, Performance over the course of task. Training phase: blocks 1-3. Testing phase: blocks 4-9. Shaded region indicates ± 2 s.e.m. **c**, Variability in performance across subjects. Plotted are distributions of performance at three task phases: late training phase (block 3), early testing phase (blocks 4-6), and late testing phase (blocks 7-9). Each distribution is shown as a violin plot (Gaussian kernel, s.d. = 0.3) and box plot (median, IQR, and range). **e**, Behavioral patterns across subjects. Data from early testing (blocks 4-6); behaviors and plotting conventions follow those in Fig. 3. Results are presented in two rows (top row: performance (proportion correct); bottom row: RT (proportion of maximum value (Max; 522 ms) in trial type matrix (leftmost plot); in sec otherwise)). Column 1: averages across subjects by trial type. Columns 2-4: mean ± 3 s.e.m. across subjects. Trial types follow those in panel Fig. 3a (column 2: symbolic distance; column 3: end item; column 4: “end order”), in addition to distinguishing between choice 1 vs. choice 2 trial types (red vs. blue, respectively). **f**, End order behavior in four example subjects. Each plot shows RTs (in sec) from an individual subject (colored circles: individual trials; dark circle and error bars: mean ± s.d.), separately plotting trials in which the end item (A or G) occurred either 1st or 2nd. Of 292 total subjects, 189 (65%) vs. 103 (35%) showed the 1st vs. 2nd faster pattern; of these subjects, 48% (1st-faster) vs. 6% (2nd-faster) showed significance at the p < 0.01 level (RTs of 1st vs. 2nd trials, rank-sum tests). **g**, Histogram of end-order behavior across individual subjects. The behavior was quantified as the difference of average RTs divided by their sum (end order index; average RTs calculated for trials where end items (A and G) occurred 1st vs. 2nd). All data from n = 292 subjects, with panels e-g presenting data from early testing (blocks 4-6).

To assess the task paradigm, we first evaluated whether subjects successfully performed inference. We analyzed two groups of subjects: (i) those showing above chance (>50%) performance on training trials in the final training phase block (292 of 392 (74%) subjects), the minimal degree of task ability prerequisite to testing inference, and (ii) those proficient in training trials (>80% performance in final training phase block; 65 of 392 (17%) subjects), similar to previous studies that initially establish TI task paradigms [51, 115]. On the first presentation of ‘critical’ trial types (testing trials not containing end items; e.g. B vs. D), either group of subjects performed well above chance: for (i), 60% ± 1.2% (s.e.m.) (vs. 50%, p < 10*^−^*^14^, signed-rank); for (ii), 75% ± 2.4% (vs. 50%, p < 10*^−^*^9^, signed-rank); subsequent analyses were carried out in the more inclusive group (i), as previous work on TI indicates that behavioral patterns can manifest even when subjects show relatively low performance on training trials (e.g. [116]). Across the testing phase, these subjects performed consistently above chance (65-85% mean performance, **Fig. 8c**), while also showing considerable variability in performance (**Fig. 8d**), as is generally observed in TI studies (e.g. [117]).

Importantly, subjects doing delay TI showed the classical behavioral patterns seen in previous TI studies, a result that was apparent when trial types were considered (**Fig. 8e** and **Fig. S4**). In particular, both symbolic distance and end-item effects were expressed, each in both performance and RTs (symbolic distance: **Fig. 8e**, 2nd column; end-item: **Fig. 8e**, 3rd column).

The above findings – indicating both successful inference and classical TI behaviors – introduce delay TI as a viable experimental paradigm, and, further, suggest that subjects’ performance of delay TI may share an underlying basis with that of traditional TI task paradigms.

We lastly found that subjects widely expressed the end order effect, the novel behavioral prediction of neural models performing delay TI (**Fig. 4**). The effect was initially observable in plots of RTs by individual trial types (**Fig. 8e**, trial-type matrix of RTs with lower values in first and last rows; **Fig. S4b** and **S4d-f**, manifest as an “X” pattern in average RTs), both of which indicated specifically that subjects responded faster when the end item (A or G) was the 1st item (1st-faster version); plots of RTs averaged across trial types (**Fig. 8e**, last column) and RTs in individual subjects (**Fig. 8f**, additional quantification in caption) confirmed the effect. Lastly, as with separate instances of neural models (**Fig. 4**), we measured the end order effect across individual subjects using a quantitative index, finding that 1st-faster behavior occurred on a distribution that was wide yet skewed to values <0 (vs. 0, p < 10*^−^*^24^, signed-rank) (**Fig. 8g**). Strikingly, this distribution was overtly inconsistent with that of simple-regime f-RNNs, which consistently show values >0 (**Fig. 4**), with quantitative comparison of index values underscoring the mismatch (**Fig. S9**). This result furthermore suggests that the solution expressed by simple-regime f-RNNs (simple solution; **Fig. 5** and **Supplementary Diagram**), while intuitive, may not accurately reflect the underlying system responsible for task performance.

## 10 Discussion

In this study, we generated, analyzed, and experimentally tested a collection of neural models of transitive inference (TI), a classical cognitive task that distills relational inference into a simple yet essential form. Motivated by the naturalistic and potentially intrinsic interrelationship between relational inference and working memory (WM) [74–77], our study introduces a new task paradigm – “delay TI” – that imposes an explicit WM delay between presented items. We found that trained recurrent neural networks (RNNs) not only performed delay TI, i.e. generalized to all novel combinations of inputs (**Fig. 2**), but also expressed behavioral patterns long documented in living subjects performing TI (**Fig. 3**). Investigating delay TI also disclosed a previously undescribed order-dependent behavior, which we termed the “end order” effect, which was expressed in RNNs in either of two distinct versions (1st- vs. 2nd-faster versions, **Fig. 4**).

We subsequently identified a neural solution to delay TI characterized by simple collective dynamics and geometry (population-level “subtraction,” **Fig. 5** and **Supplementary Diagram**). This solution, which was expressed in a subset of RNNs optimized for efficiency and having modifiable feedforward connectivity (simple-regime f-RNNs), led us to identify a set of testable activity predictions unique to these models, among others that successfully performed the task (**Fig. 6**, **7**; summary of main predictions in **Table 3**).

Lastly, in a large-scale experimental study of delay TI, we found that human subjects successfully performed the task and also showed classic behavioral patterns seen in traditional TI tasks (**Fig. 8**). Strikingly, subjects expressed the end order effect, doing so in mainly one of two alternative versions (1st- rather than 2nd-faster), thus providing grounds for ruling out neural models uniformly expressing the alternative version (2nd-faster).

Prior work on relational inference in the brain has often focused on task paradigms that rely on stimuli that are challenging to isolate (e.g. spatial tasks [21,24,47,118,119]), test multiple relations at once (e.g. tasks with linguistic responses and/or episodic elaboration, e.g. [120–123]), or do not require behavioral report of inference. Possibly as a result, there are relatively few hypotheses and available models that clarify or explain how neural systems accomplish relational inference (generalize in accordance with a relation) at the explanatory levels of population-level neural activity and behavior. In the present study, we developed a task paradigm (delay TI) and a neural approach (task-trained neural networks) suited to meet these challenges. It is also worth emphasizing that the TI paradigm presently studied is implicit, i.e. does not provide semantic or isolated perceptual cues regarding the underlying transitive relation, thereby minimizing the role of linguistic ability and affording a bridge to the extensive literature on TI in animals [53, 58].

We initially found that RNNs trained via standard optimization procedures commonly performed TI perfectly (transitive generalization; **Fig. 2**, **Tables 2** and **5**). By itself, this finding is novel as it is not generally known whether largely unstructured learning models, which trained RNNs and other NNs exemplify, implement the inductive bias required for transitive generalization (examples in feedforward models in **Fig. S2** and [104, 105]). Interestingly, this finding raises the question of what exact components of learning systems implement such ‘relational’ inductive biases [5], for which transitivity is an archetype. Indeed, in TI, existing reinforcement learning (RL) models, which represent the most behaviorally relevant models to date [60], have sidestepped this question. On the one hand, RL models that cannot acquire internal representations fail to perform TI or show behavior that is qualitatively non-naturalistic, thus disqualifying these models (e.g. Q-learning and value transfer models [60]). On the other hand, other RL models used to study TI have been pre-configured to have transitive or otherwise ordered internal representations (e.g. a score or set of ordered lists [60]). By contrast, our observation that TI was often expressed in trained RNNs, learning systems that are not pre-configured as such, invokes the possibility that relational inference can emerge in a wide range of learning systems, moreover by virtue of principles not yet identified. We highlight this matter as an important direction for future work: in our investigation, we subsequently focused on how models generalized after having acquired the ability to respond correctly on training trials (AB, BC, CD, etc.).

Crucial to our approach was to identify, where possible, multiple solutions to TI. To do so, we investigated whether and how RNNs performed delay TI when two neurobiologically relevant factors varied — learnable connectivity (i.e. fully-trainable RNNs (f-RNNs) vs. recurrent-trainable (r-RNNs)) and training regime (regularization and initial synaptic strength; ‘simple’ vs. ‘complex’ training regimes) (**Fig. 1d**, **Table 1**). We found that each of these RNN variants could perform TI, and moreover identified a set of behavioral and neural predictions that distinguish between four representative variants (**Table 3**). Importantly, RNN variants expressed qualitatively different versions of the end order behavior (**Fig. 4**, **Table 3**, second column), an indication that networks adopted different solutions to the task.

At the same time, the neural activity expressed in the various networks performing TI suggested a common dynamical principle: namely, rotation. This was initially suggested by the “subtractive” solution (in simple-regime f-RNNs, **Fig. 5e**), in which a single oscillation rotates activity states in activity space during the delay. Beyond this solution, we observed that a single oscillation was also sufficient dynamics to perform TI even in networks that could not express an orderly arrangement of activity states from feedforward input (due to lack of modifiable feedforward connectivity; simple-regime r-RNNs, compare **Fig. 5** with **S6**); this indicates that a single rotational transformation can be used to perform delay TI for additional activity geometries elicited by feedforward input. More broadly, a single oscillation could be a simple case of a wider set of dynamical patterns enabling transitive generalization, with rotation as the essential operation. The idea of generalized rotational transformations also led us to identify an angle-based activity pattern that distinguished all RNN variants in accordance with task behavior (mean angle change, **Fig. 6c**; behavior prediction in **S8b**).

Geometric analyses of activity (**Fig. 6**), in addition to the ability of RNNs lacking modifiable feedforward input (r-RNNs) to perform the task, indicated that there exist a variety of activity geometries capable of supporting transitive generalization. Intriguingly, complex-regime RNNs often showed a “V”-shaped arrangement of activity states (manifesting at the end of the delay and differing from the more strictly collinear geometry seen in simple-regime f-RNNs; examples in **Figs. 6a** and **S7a**; see also **S7c**), similar to that reported in recent studies testing human subjects on a transitive hierarchy of items [105,114]. The activity pattern is also notable in that it typically exhibits a segregation of the end items (A and G) from the middle items – this segregation echoes the finding that, in these same networks, the encoding of end items is exceptionally amplified during the delay (invariably seen in complex-regime RNNs; ‘1st-dominant’ encoding, **Fig. 7**). With respect to function, this encoding strategy is advantageous as it yields faster responses in trials where item 1 is an end item, i.e. when item 1 sufficiently indicates the correct response prior to and irrespective of item 2; in parallel, among RNN variants, complex-regime networks showed the most robustness to changes in delay length (**Fig. S3b**). These results highlight an unexpected connection between (complex-regime) RNNs and experimentally observed activity geometry in related tasks. More broadly, these results raise the possibility that the observed “V” activity geometry was generated by learned recurrent dynamics (rather than particular input encodings or other mechanisms [105]), and, further, suggest specific advantages for behavior.

More directly, our experimental study of delay TI in human subjects provided empirical grounds for discriminating between neural models. Besides finding that subjects showed well-established TI behaviors (**Fig. 8e**), we found that subjects showed the 1st-faster version of the end order behavior, both individually and in aggregate (**Fig. 8f-e**). This finding excludes simple-regime f-RNNs, as these models consistently show the 2nd-faster behavior, and instead suggests that neural models expressing the 1st-faster version may be more accurate. RNNs expressing the 1st-faster version (**Table 3**) have three unorthodox properties worth highlighting: (1) non-learnable feedforward connectivity (r-RNNs), (2) relatively larger scale of initial connectivity weights (intermediate- and complex-regime), and (3) weaker or no regularization (intermediate- and complex-regime). (It is also worth noting that, amongst RNN variants, intermediate-regime f-RNNs provided the best quantitative match to human data (**Fig. S9**).) Though we do not here interpret RNN training as a model of task learning in the brain, these three properties may nonetheless be understood as high-level constraints on the neural system in the brain responsible for acquiring and expressing delay TI, and potentially relational abilities more generally.

These properties invoke several further considerations regarding underlying neurobiology and behavior. Property (1) implies that the neural system in the brain responsible for performance of the task may rely upon learned recurrent dynamics rather than on learned feedforward input — two fundamentally different neural implementations of the underlying relational ability. Indeed much prior work on neural substrates of relational abilities (whether construed as schemas, relational memory, semantic knowledge, etc.) has focused on feedforward operations, despite the fact that the relevant neural systems are invariably extremely recurrent. In addition, the empirically suggested plausibility of properties (2) and (3) is unusual in that these properties run counter to prior work indicating that task-trained RNNs best matching neural responses in the brain have the opposite properties, i.e. relatively smaller scale of initial connectivity weights and strong regularization [48, 95, 100, 124]. Lastly, property (1) and properties (2) and (3) yield RNNs that show the 1st-faster end order behavior, which, as described above, is behaviorally advantageous.

Further experimental data is required to distinguish between neural models. Importantly, four representative RNN variants in our study were distinguished by neural activity predictions (at the population level; summarized in **Table 3**); it is thus in principle possible to discriminate between these models with experimentally collected neural data. With respect to brain regions, any number of brain structures receiving appropriate sensory input and capable of supporting WM and the learning of training trials are relevant; TI has moreover been observed in a wide variety of animal species [53, 58]. In the mammalian brain, the leading candidates are brain regions linked to relational inference and/or WM, including prefrontal cortex (PFC) and hippocampus [125–131], both known to be required for TI performance [61, 132]. Notably, discriminating between neural models of delay TI has the potential to provide insight beyond how these brain regions enable TI, but also how these brain regions contribute to performance of other cognitive tasks. Such insights could arise both from the structure of specific solutions and from identifying high-level constraints (e.g. learnable connectivity).

The findings in the present study may also relate to the neural basis of other cognitive functions. Of direct interest are neural activity patterns relevant to abstraction — whether at the level of single cells (e.g. place and grid cell firing [14,55] and other firing having abstract correlates [133,134]) or neural populations (e.g. activity geometries [45, 100, 135–137], dimensionalities [138–141], and re-activation patterns [142–145] suitable for generalization). In our approach to TI, we deliberately chose not to seek to fit or capture these neural activity patterns, instead stipulating relatively unconstrained neural models. Indeed there may exist important relationships between such activity patterns and those expressed in the neural models here presented. It is also worth emphasizing that our findings do not directly address learning processes, for which prior studies have proposed various models and mechanisms [60, 81, 146–150] (including for explicit variants of TI, where human subjects are informed of the transitive hierarchy [105, 151, 152]). Further, our analyses and neural activity predictions focus on delay period activity, leaving open the question of whether and how neural activity following presentation of both items may contribute to transitive generalization. More broadly, it is worth pointing out that there exist any number of other types of relational inferences (e.g. spatial navigation), tasks, and scenarios that incorporate considerably higher, and potentially important, complexity. In conjunction, in the brain, the relevant regions linked to TI and relational inference also support or pertain to cognitive capacities such as structure learning, episodic and semantic memory, and imagination. This convergence of diverse cognitive functions indicates that, toward understanding their biological basis, there is a major need to synthesize approaches.

## Acknowledgements

We thank S. Lippl, J. Johnston, L. Tian, M. Triplett, C. Monfredo, K. Kay, B. Antin, J. Cunningham, and members of the Columbia Center for Theoretical Neuroscience for comments and discussion, and R. Yang for guidance on model training. This work was supported by the the Simons Collaboration for the Global Brain (521921 and 542981), NIH grants (R01 MH111703, V.F.), the NSF NeuroNex award (DBI-1707398), the Gatsby Charitable Foundation, and an NIMH K99 (MH126158-01A1, KK).

## Author contributions

K.K. conceived study in discussion with X.W., V.F., and L.F.A. All authors contributed to study design. K.K. implemented models and performed all analyses. N.B. implemented human study and collected data. K.K. wrote manuscript with input from all authors.

## Code availability

Code and trained networks will be made available upon publication.

## Methods

### Task

Transitive inference (TI) is a classic cognitive task that requires subjects to infer an abstract ordered relation – here, transitivity (**Fig. 1a**) – between items not previously observed together, i.e. using A > B and B > C to infer A > C (**Fig. 1b**). TI defines test cases expressly as novel recombinations of training inputs, thus primarily testing relational rather than statistical inference. We focused on a 7-item version of TI, in which there are 12 training trials and 30 test trials (training: A vs. B, B vs. C, etc.; test: A vs. C, B vs. D, etc.; see **Fig. 1b** for diagram of trial types and correct responses), though in pilot work we found that our approach could be generalized to fewer or more items with qualitatively similar results.

Given the interrelationship between relational inference (exemplified by TI) and working memory (WM) [75,76], we investigated TI in a task format that explicitly imposes a delay that necessitates WM (**Fig. S1a**, delay format). Further, for comparison to previous modeling work [60,104] and for potential insight, we also studied the traditional task format, for which the presentation of task items (A, B, C, etc.) is simultaneous (traditional TI, diagram in **Fig. 1c**). It is worth noting that in some prior TI studies, a delay between stimuli is implicit in the free-exploration afforded to subjects (e.g. [52]).

Besides WM, an important difference between the delay vs. traditional format is that the traditional format requires twice as many input parameters as the delay version (e.g. twice as many input connections in a neural system). This difference may make the delay version not only more difficult to perform, but also more neurobiologically accurate with respect to neural systems underlying abstract cognition: these systems look to receive extremely diverse inputs, implying that the extent of input connectivity is relatively constrained [153].

### Input stimuli (items)

A panel of input stimuli corresponding to items A, B, C, etc. was constructed for each model instance. In TI in living subjects, an item is an arbitrary sensory object (e.g. image, odor) with no features that are significant *a priori*. To capture this property, items were represented as randomly generated input vectors **u** (**u***^A^*, **u***^B^*, etc.), modeling sensory-driven activity in upstream neurons. For simplicity, each **u** was drawn from a multivariate standard normal distribution of dimension *N^in^*.

The identity of items 1 and 2 varied by trial type (e.g. AB, BA, AC, BC, etc.; see **Fig. 1b** for all trial types). *N^in^*was chosen to be 100, matching the size of the hidden layer of the neural models (MLP and RNN), which was set to *N* = 100 (see below). This ensured that input corresponding to each item presentation elicited patterns of activation that would be uncorrelated in the activity space of the neural models (at least prior to training) thereby simulating arbitrary sensory stimuli. Note that TI (and relational inference more generally) is not defined in terms of particular stimulus features, and indeed is most rigorously tested in the absence of any stimulus features indicating items’ rank in the transitive hierarchy [154].

### Model architectures

Three model architectures were studied: a recurrent neural network (RNN), logistic regression (LR), and a multi-layer perceptron (MLP). Each was implemented in Python using the NumPy and PyTorch [155] packages, in addition to custom code for RNNs and all subsequent analyses.

#### Recurrent neural network (RNN)

To investigate population-level neural dynamics, we studied the standard continuous-time RNN:

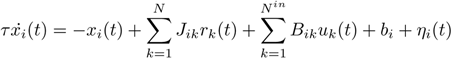

where *x_i_* are activity of the recurrent units, *r_i_* are the corresponding rates, *N* is the number of recurrent units (100 for all networks), *N^in^* is the number of input units, and *τ* is the time constant. The rates *r_i_* derive from the activations *x_i_* via a tanh nonlinearity, *r_i_*= tanh(*x_i_*).

The tanh non-linearity was chosen because we found it to be the most effective for generating and analyzing network dynamics; further, in pilot work we found that other non-linearities (e.g. rectified tanh) yielded RNNs that exhibited unrealistic behavioral patterns when simulated (see below for description of behavioral simulations; rectified-tanh RNNs exhibited an unrealistic response bias favoring choice 1 over 2, or *vice versa*). We note that RNNs in the present study are intended to model neural activity (and generate testable predictions) at the neural population level.

The network units interact via the recurrent synaptic weight matrix **J**. The input to the system is **u**, the activity of the set of *N^in^* input units that influence the network through input weights **B**. The output of the system is **z**, the activity of a set of *N^out^*output units, each defined to be a linear readout of activity in the recurrent units:

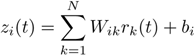

Each output unit *z_i_* is a weighted sum of network rates with weights, **W_i_**, with a constant bias, *b_i_.* In all models, three output units were implemented (*N^out^* = 3), corresponding to three alternative behavioral actions (see below, Model outputs). All analyses of neural activity were performed on *x_i_*; analyses of *r_i_*yielded similar predictions.

Network simulations were performed using Euler’s method with discrete time step Δ*t* = *τ/*10. Intrinsic single-unit noise *η_i_*(*t*) was generated at each time step by drawing values from a Gaussian random variable with zero mean and s.d. of 0.2.

Prior to training, the parameters of the model were initialized as follows. The entries of **J** were initialized as draws from a normal distribution with zero mean and variance 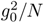. The entries of **B** were initialized as draws from a normal distribution with zero mean and variance 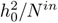. The elements of **W**, and all bias terms, were initialized to 0. Both *h*_0_ and *g*_0_ were hyperparameters that were systematically varied across RNNs (see **Table 5** and RNN variants below).

#### Logistic regression (LR)

The LR model (schematic in **Fig. S2a**) was studied to clarify the possible role of feed-forward connectivity in performing TI. Each LR consisted of two linear readouts (corresponding to choice 1 and 2) each of which had coefficients for every input dimension *N^in^* for each of the two items presented. In each simulation of the model, Gaussian noise (zero mean, s.d. of 0.2) was added to each of the readouts.

#### Multi-layer perceptron (MLP)

In addition to the LR model, single-layer MLPs (schematic in **Fig. S2a**; N = 100 hidden units, fully connected) were studied to clarify the possible role of feedforward connectivity in TI. Entries of the input weight matrix were initialized as draws from a normal distribution with 0 mean and variance 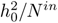, with *h*_0_ = 1; entries of the output weight matrix were initialized to 0; all biases were initialized to 0. In each simulation of the model, Gaussian noise (zero mean, s.d. of 0.2) was added to each of the hidden units. Relevant results in MLPs are also reported in two previous studies ([104] for MLPs with three hidden units solving five-item TI, and [105] for MLPs constrained to have symmetric input weights).

### Model input

#### RNN

The input to the RNN in trial *m*, **u**(*t, m*), consisted of the presentation of items 1 and 2 with an intervening delay, dividing three periods in each trial: rest, delay, and choice (**Fig. S1a**). Item presentations were modeled as instantaneous pulses (one timestep), as TI (and relational inference more generally) is not dependent on sensory input of a particular duration.

RNNs were trained on one of three input formats in which the duration of the delay period differed (delay variants): (i) basic, (ii) extended, and (iii) variable. In (i), the rest, delay, and choice periods lasted 0.5*τ*, 2*τ*, and 2*τ*, respectively. A delay duration of 2*τ* was sufficiently long to yield different neural implementations of TI in trained networks, and is the minimal delay duration relevant to working memory [156, 157]. In (ii), the trial periods lasted 0.5*τ*, 6*τ*, and 6*τ*, respectively. In (iii), the rest period and total trial duration were the same as (ii), but the delay duration was varied randomly from 2*τ* to 6*τ* (uniformly across individual trials by shifting the time of item 2 presentation); subsequent testing and simulations of RNNs trained on (iii) were performed using the maximum delay duration (6*τ*). All delay variants yielded RNNs that performed TI (**Table 4**) and made qualitatively similar behavioral predictions (**Fig. S3c-d**); unless indicated otherwise, results from RNNs trained on (i) are presented throughout the study.

**Table 4:**
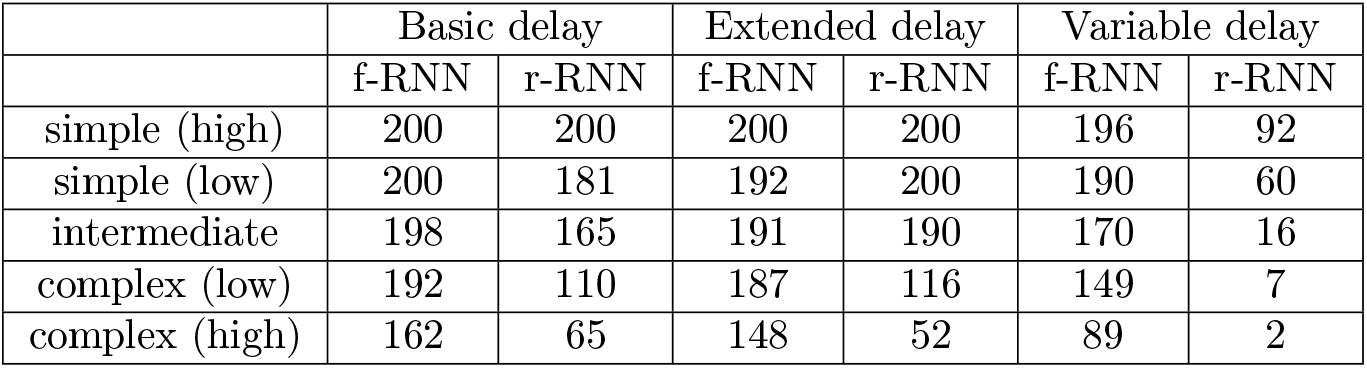
Number of RNNs that fully generalized out of 200 trained instances. All trained instances responded correctly on all training trials, regardless of generalization performance.

#### MLP and LR

The input to the feedforward models (LR and MLP) in trial *m*, **u**(*m*), consisted of the joint (simultaneous) presentation of items 1 and 2, requiring twice the input dimensionality of RNNs (i.e. given a fixed dimensionality for individual input stimuli, i.e. items); thus the feedforward models had *N^in^* = 200 rather than *N^in^* = 100 as in the RNNs (diagrammed in **Fig. S2a**).

### Model output

#### RNN

The output from the RNN was composed of three output units *z*_1_, *z*_2_, *z*_3_ corresponding to three behaviors: choice 1, choice 2, and rest, respectively. In training, the target output 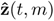 was defined for every time point *t* and for each trial type *m* such that the correct output unit was activated above resting levels during the choice period (target values: resting: 0, activated: 5; diagrammed in **Fig. S1b**).

In example outputs (**Fig. 2**, top row, and **Fig. S3a**), the choice value plotted was the difference in the readout values for *z*_1_ and *z*_2_ averaged over the last half of the choice period and normalized to the magnitude of the largest such difference value across all trial types.

#### RNN behavior

The behavior (i.e. the choice response and response time (RT)) of an RNN in a trial was defined using an established criterion [106]. The *z*_1_ (choice 1) and *z*_2_ (choice 2) output units were passed through a simple monotonic saturating function ranging in value from 0 to 1:

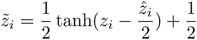

where 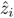 is the target value of the output unit.

The **response** (choice in trial) was defined by the identity of the output unit (choice 1 vs. choice 2, see above) that first reached a fixed threshold value of 85% in the choice period. Under certain conditions (i.e. when additional noise was added to RNNs (see further below), or cases when the RNN was presented with same-item stimuli, e.g. AA, BB, CC, etc., **Fig. 2**; these trial types were not evaluated in this study), the threshold was not reached for output unit. These trials are shown in plots as ‘no response’ trials (**Fig. 2**).

The **RT** was defined as the time of the response, measured as the time elapsed from *t*_2_ (the time of presentation of item 2), normalized to the maximum duration of the choice period (0 to 1). In a subset of plots, the RT was normalized to the maximum RT observed across trial types (e.g. trial-type matrices of RTs in the first column of **Fig. 3**), and described accordingly.

#### MLP and LR

For both feedforward models, the output was composed of two output units *z*_1_, *z*_2_, corresponding to choice 1 and choice 2, respectively. In training, the target output 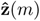 was defined for each trial type *m* such that the correct output unit for each trial type (choice 1 vs. 2, **Fig. 1b**) was activated (value for active: 1, value for not active: 0). The response for a given trial was defined by the identity of the output unit which had the higher activity value. In example outputs (**Fig. S2b**), the choice value plotted was the difference in the readout values for *z*_1_ and *z*_2_ normalized to the magnitude of the largest such difference value across all trial types.

### Model training

Models were optimized (trained) solely on training trials and not test (inference) trials. The ability of trained models to respond correctly to inference trials thereby mirrors that of living subjects that have only experienced or learned from training trials. In this way, analysis of models that respond correctly on test trials (i.e. perform inference) can be studied to identify putative neural implementations. Parameter updates were performed for batches of training trials, where each batch consisted of 128 trials randomly sampled from the training trial types defined by the task (diagrammed in **Fig. 1b**).

#### RNN

RNNs were trained to minimize *E_task_*, the average squared difference between **z**(*t, m*), the readout of the network on trial m, and 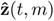, the target output for that trial:

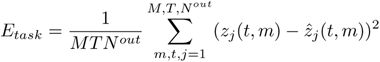

where m corresponds to different training trials, T corresponds to the length of the trial (in time steps), and *N^out^* is the number of readout units. *E_task_* stipulates that the optimization procedure generate networks that respond correctly in training trials.

The overall error function *E* was comprised of *E_task_*and two additional terms that implement regularization, which has been found to promote neurobiologically accurate solutions in trained RNNs [48, 95, 96]. The two terms were *R_L_*_2_, a standard L2 regularization on input and output synaptic weights, and *R_F R_,* a regularization on the network rates.

The overall error function was

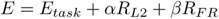

where the *α* and *β* hyperparameters set the strength of each type of regularization.

The first regularization term is a standard L2 penalty on input and output synaptic weights:

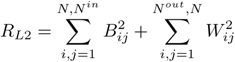

The second regularization term is a metabolic penalty on rates in the network:

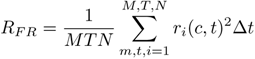

Both terms have been found to promote neurobiologically realistic responses in trained RNNs (e.g. [48, 95, 96]).

The objective of training was to minimize E by modifying the network parameters **J**, **B**, **W**, **x**(t=0), and constant bias terms.

Training was implemented with the Adam optimizer [158], with updates to the network parameters calculated using backpropagation through time [159, 160]. Parameter updates were performed for training sets (batches) of 128 trials, where the trials in each batch were randomly sampled from the task-defined training trials (diagrammed in **Fig. 1b**). Training was stopped when the RNN responded correctly on all trial types (training and test; see above for response criteria) in the absence of noise (*η_i_* set to 0), or when *E_task_*fell below 0.1; for RNNs trained on the variable delay input format, correct responses were further required when the time of item 2 was advanced earlier in time by 75% of the longest delay duration. Up to 30,000 training epochs were run.

#### MLP and LR

Each feedforward model was trained using the Adam optimizer with cross-entropy loss. For the LR models, training was performed to convergence (i.e. until loss did not improve for 1000 training epochs); for MLPs, training was performed until the network responded correctly on all trial types (training and test; see above for response criteria) in the absence of noise (*η_i_* set to 0). For both models, a weight-decay term (given by the L2-norm of all parameters) scaled by hyperparameter *α* was included in the loss function, with similar results across a range of values (presented are 0.1 for LR and 0.001 for MLP).

### RNN variants

To identify multiple biologically relevant neural implementations of TI, two classes of RNN variants were studied:

First, RNN variants that differed in learnable connectivity: f-RNN and r-RNN. f-RNNs (fully-trainable RNNs) were RNNs where all connection weights (feedforward and recurrent) were trainable; r-RNNs (recurrent-trainable RNNs) were RNNs for which feedforward weights (**B** and **W**, see above) were not allowed to be modified from their initial random values (Gaussian draws) in training. In addition, we separately found that RNNs for which input feedforward input weights (**B**) were not allowed to change showed similar results to r-RNNs. Since these latter RNNs were a relatively closer point of comparison to f-RNNs, these models were analyzed and presented in **Fig. 5** and **Fig. S5a** (last row).

Second, RNN variants that differed with respect to initial connectivity strength (prior to training) and regularization (during training) – jointly termed “training regime” and defined for five classes: simple (high), simple (low), intermediate, complex (low), and complex (high) (parameter values in **Table 5**). The ranges of parameter values are comparable to those in prior work using trained NNs to model other tasks (e.g. [48, 95, 100]). In pilot studies, parameter values beyond the ranges presently used tended to yield models that showed unrealistic behavior, e.g. failures to generalize.

**Table 5:** Training regime parameters. *h*_0_, input gain; *g*_0_, recurrent gain; *α*, weight regularization; *β*, metabolic regularization.

| | $h_0$ | $g_0$ | $\alpha$ | $\beta$ |
| --- | --- | --- | --- | --- |
| simple (high) | 1.0 | 0.5 | 1.0 | 1.0 |
| simple (low) | 1.0 | 0.5 | 0.01 | 0.01 |
| intermediate | 1.0 | 1.0 | 0.01 | 0.01 |
| complex (low) | 1.0 | 2.0 | 0 | 0 |
| complex (high) | 1.5 | 4.0 | 0 | 0 |

In reporting results, we generally use the terms “simple-regime” and “complex-regime” to refer to the simple (high) and complex (high) training regimes, respectively, though in some cases we use “simple-regime” to refer to both simple (high) and simple (low) (and similarly for “complex-regime”). If the case, this is made explicit where presented. Ten RNN variants were studied: two types of learnable connectivity variants (f-RNN and r-RNN) by five types of training regime variants.

In addition to these two types of variants, for each RNN variant (e.g. simple-regime f-RNN) a collection of individual instances (different random initializations) were studied. In particular, we trained 200 instances of each RNN variant, subsequently studying only those models that performed TI perfectly (correct responses to all trial types) under noise-free conditions (*η_i_*set to 0). This subset of instances were then subject to behavioral and neural analyses.

### Behavior simulation and behavioral patterns

To investigate behavioral patterns across models (**Fig. 3, 4, 7, S2, S3, S4, S8**), models were simulated on all 42 trial types (12 training and 30 test, **Fig. 1b**). To simulate average levels of performance that were realistic to living subjects performing TI and showing characteristic behavioral patterns (>50% performance in training trials and <100% performance on trial types with large symbolic distance; see example monkey data in **Fig. S4a**), we took the following approach.

#### Simulation approach

For each model instance that performed perfectly (correct responses on all trial types) under noise-free conditions, we added progressively larger amounts of intrinsic noise to model units (s.d. of *η_i_*, increased from 0 to 3.0 in increments of 0.05; LR: readout units, MLP: hidden-layer units, RNN: hidden-layer (recurrent) units) until the performance of the model (averaged over 500 simulations across all trial types) satisfied these basic performance criteria: >50% training performance for both choice 1 and 2 training trials and <96% performance on the largest symbolic distance trials (AG and GA). All RNN instances meeting these basic performance criteria were subsequently analyzed; for subsequent behavioral analysis, 500 simulations (of all trial types) at the identified noise level were run.

With the addition of noise, a subset of simulated trials (~10%) did not meet the output activity threshold criterion (fixed at 85%) for a response (no response trials, see above). Though it is possible to use additional response criteria to estimate network responses in these trials [140], the following approach was taken for simplicity of interpretation: choice in these trials was randomly designated as either 1 or 2, and these trials were excluded from analyses of RT.

#### Symbolic distance effect

Trial types differing by the magnitude of the difference in rank between items (i.e. distance 1: AB, BA, BC, CB, CD, DC, DE, ED, EF, FE, FG, GF; 2: AC, CA, BD, DB, CE, EC, DF, FD, EG, GE; 3: AD, DA, BE, EB, CG, GC; 4: AE, EA, BF, FB, CG, GC, 5: AF, FA, BG, GB, 6: AG, GA). Schematic of trial types in Fig. 3a (second column).

#### End item effect

Trial types differing by whether or not they contain end items A or G. Schematic of trial types in Fig. 3a (third column).

“End order” effect

Trial types for which end items (A or G) were presented 1st (item 1) vs. 2nd (item 2). Schematic of trial types in Fig. 3a, last column. Note that trial types containing both end items (i.e. AG and GA) were not included. The end order effect refers the (sequential) order of item presentation, and is therefore specific to delay TI.

The end order index (**Figs. 4**, **8**) was defined as:

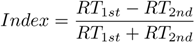

where *RT*_1*st*_ is the average RT over trials where end items were presented first (item 1), and *RT*_2*nd*_ is the average RT over trials where end items were presented second (item 2).

### Visualization of population activity

To clarify the implementation of TI in neural models, population-level activity was visualized by performing PCA and plotting activity in the top PCs. For each model, PCA was performed for two sets of activity: (1) a comprehensive set (presented in **Figs. 5** and **S6**), comprised of activity across all time points and trial types (all 12 training and 30 testing types; % variance explained in **Table 3**) or (2) at the end of the delay (presented in **Figs. 6** and **S7a**), comprised of activity from the final time step of the delay period (7 trial types corresponding to the rank of item 1, i.e. A, B, C, etc.). Activity set (2) was visualized to clarify geometric relationships between trial types in the delay, when differences in activity are solely determined by the identity of item 1 (i.e. rank of item 1). All activity vectors were taken under noise-free simulations (*η_i_*set to 0).

### Activity axes

Visualization of population activity in RNNs performing TI (**Figs. 5**, **6**) suggested that the underlying neural implementations were characterized by specific arrangements (in activity space) of activity states with respect to two directions (axes), each defined on the basis of different trial types in the task (visual schematic of each axis in **Fig. S8a**).

The first was the **choice axis**, defined as the direction pointing from the activity states of Choice 1 vs. Choice 2 trial types (21 trial types each; red vs. blue, respectively, in **Fig. 1b**) during the choice period (period following the delay, **Fig. S1a**). The choice axis was calculated as a unit-normalized vector pointing from the mean of activity vectors across Choice 1 trial types to the mean of activity vectors in Choice 2 trial types, both for activity averaged over the first quarter of the choice period.

The **cross-condition mean** (**XCM**; yellow line in **Figs. 5a**, **5b**; [111–113]), was defined and calculated as the average trajectory across all trial types. The XCM was moreover calculated in two ways: for visualizations, the XCM was calculated for every time point (**Fig. 5a, 5b**); for quantifications, the XCM was calculated by taking the mean neural activity across a given time window (e.g. first or last quarter of the delay period). The **XCM axis**, defined as the direction pointing from the activity states at the beginning of the delay period to the end of the delay period, was calculated as a unit-normalized vector pointing from the XCM of the first quarter of the delay period to the XCM of the last quarter of the delay period.

The **readout axis** was defined as the set of weights from the recurrent units to the choice 1 output unit (**W_1_**, see above).

### Inference of linear dynamics

To identify dynamical components expressed in RNNs performing TI, we fit neural activity from the delay period to an unconstrained linear dynamics model (Ẋ = *XA*; least-squares fit) from noise-free simulations (*η_i_* set to 0) across all time points during the delay and across all trial types. In the delay period there are 7 trial types corresponding to each possible item 1 (A through G). The fit was performed for the top 10 PCs of delay-period activity. *R*^2^ values of the fit were relatively high (0.5-0.9; see **Fig. S7**, first column, for values across networks). Eigenvalues of the A matrix were subsequently plotted for each type of RNN variant (**Fig. S5a**, columns 2-6).

### Fixed point analysis and linearization

To identify dynamical components of RNNs performing TI, we used fixed-point analysis and linear approximation methods [97, 108, 109].

Fixed-point finding was implemented using custom code in PyTorch, following an established method [97, 108]. Optimization via gradient descent (Adam optimizer) was used to identify activity states in which the speed of RNN dynamics was minimized (mean-squared error loss). The optimization was seeded using activity states from noise-free simulations of trial types in the TI task (specifically at these time points: 0, time of item 1 presentation, time of item 2 presentation, halfway through delay, the last time point, and the last timestep when trials were simulated with 100 additional timesteps), in addition to 5 batches of 50 activity state seeds, each of which were drawn randomly from activity states of trials in which item 1 and item 2 were randomly jittered in time across the trial. Each batch was optimized for 50000 epochs and stopped after 5000 epochs with no improvement in loss. Candidate FPs were those activity states for which the speed was lower than 10*^−^*^5^. Redundant candidate FPs were eliminated by requiring that, between candidate FPs, the activity of every recurrent unit differed by more than 10*^−^*^5^.

To obtain the Jacobian matrix A of linearization, two methods were used. The first was analytic (based on the weight matrix [108]); the second was numerical [84], using the function grad() in the PyTorch autograd library: at each fixed point, the function grad() was used to calculate the entries of A for the trained (and frozen) RNN (with no external input). Each approach yielded equivalent results.

### Oscillation of transitive comparison

To identify the putative oscillation associated with transitive comparison (comparison oscillation), for each RNN performing TI we detected the oscillatory mode (mode with eigenvalue having a non-zero imaginary component) of highest frequency (e.g. highlighted in arrowheads in **Fig. 5b** and **6b** for example networks), as this was the mode associated with transitive comparison in simple-regime RNNs (modes with frequency ~0.5 cycles/delay, **Figs. 5d, 6d**). This detection was performed from the inferred linear dynamics matrix (with eigenvalue spectra across RNNs in **Fig. S7**), as this can also be performed for experimental neural data. The 2D linear subspace (plane) of the identified oscillatory mode was defined by the eigenvectors, which were orthogonalized and unit-normalized prior to being used to visualize population activity (**Fig. 5b**, right) and to quantify model predictions regarding activity geometry (angles of task-relevant axes with respect to the oscillation, **Fig. S8**).

### Activity geometry

To quantify patterns of population-level neural activity characteristic of different neural implementations of TI, we calculated the following geometric measures (indices). These indices were calculated for delay period activity, and were based on the following groupings of task items: *S* = 7 (all items: A, B, C, D, E, F, G), *S_outer_* = 6 (all items except D: A, B, C, E, F, G), *S_high_* = 3 (high-rank items: A, B, C), *S_low_* = 3 (low-rank items: E, F, G). Note that each item defines a trial type during the delay period; thus for delay period activity there were 7 different trial types. All indices were calculated in the top 10 PCs of delay period activity; geometric indices were measured in the full activity space of networks (N-D ambient space, reduced to top 10 PCs) to avoid assumptions or biases incurred from an intermediate step of estimating activity subspaces (such as the subspace of an oscillation). Index values were compared between networks (e.g. RNN variants); for comparison to minimally structured activity, Gaussian random vectors were substituted for every activity vector (across timesteps and trial types) and index values re-calculated 1000 times.

#### Collinearity index

To measure the degree to which population neural activity is linearly arranged in activity space (schematic in **Fig. 6a**; characteristic of the “subtractive” solution, **Fig. 5f**), we defined the following index

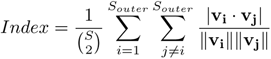

Activity state vectors (*v*) were network activity states (x) measured with respect to the cross-condition mean (XCM; yellow circle in **Fig. 6b** schematic; see also **Fig. 5a**). This index was measured at two time points: at the first timestep of delay (early; **Fig. 6b**, left column) and at the last timestep of the delay (late; **Fig. 6b**, middle column). The change over the delay (change; **Fig. 7a**, right column) was defined as late index - early index. Similar results were obtained when index values were quantified by averaging activity over time windows (i.e. first and last quarters of the delay).

#### Ordered collinearity index

To measure the degree to which population neural activity is both rank-ordered and linearly arranged in activity space (**Fig. S7b**), we defined the following index

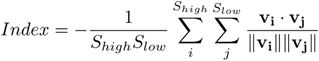

Activity state vectors (*v*) were network activity states (x) measured with respect to the cross-condition mean (XCM; yellow circle in **Fig. 7a**; see also **Fig. 5a**). Note that the negative sign results in value +1 for activity conforming to rank-ordered collinearity. This index was measured at two time points: at the first timestep of delay (early delay) and at the last timestep of the delay (late delay). The change over the delay (change) was defined as late index - early index. Similar results were obtained when index values were quantified by averaging activity over time windows (i.e. first and last quarters of the delay).

#### Mean angle

To generalize OCI change to other possible angular arrangements of neural activity, we measured mean angle change (schematic of mean angle in **Fig. 7b**), defined as

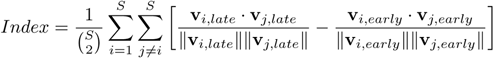

Activity state vectors (*v*) were network activity states (x) measured with respect to the cross-condition mean (XCM), and calculated for the first timestep of the delay (early) and at the last timestep of the delay (late). Similar results were obtained when index values were quantified by averaging activity over time windows (i.e. first and last quarters of the delay).

#### Mean distance

To help distinguish between angular vs. non-angular rearrangements of activity states in activity space (during the delay period), we measured mean distance change (schematic of mean distance in **Fig. 7b**), defined as

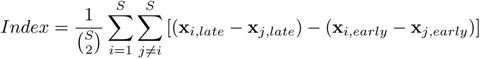

The index was calculated for network activity states (x) at the first timestep of the delay (early) and at the last timestep of the delay (late). Similar results were obtained when index values were quantified by averaging activity over time windows (i.e. first and last quarters of the delay).

#### Axis angles

To quantify angular relationships between activity patterns (**Fig. S8**), we calculated the relevant cosine angles (dot products). In particular, simple-regime RNNs (**Figs. 5** and **6**) predict two angular relationships: (1) orthogonality (cosine angle: 0) between the XCM and the plane of the oscillation associated with transitive comparison (plane of the comparison oscillation), (2) alignment (cosine angle: ± 1) between the choice axis and the plane of the comparison oscillation. The plane of the comparison oscillation was defined by the two eigenvectors of the comparison oscillation (see above, Linear dynamics), which were orthogonalized and unit-normalized, and (3) alignment between the choice axis and the readout axis. The readout axis was defined as the output weights **W_1_** corresponding to *z*_1_, the output unit correspondent with choice 1.

For (1) (**Fig. S8b**), the dot product was calculated between the XCM and each of these plane vectors. As a conservative estimate, the cosine angle was defined as the dot product having the larger magnitude.

For (2) (**Fig. S8b**), the dot product was calculated between the choice axis and each of these plane vectors. As a conservative estimate, the cosine angle was taken to be the dot product having the smaller magnitude.

For (3) (**Fig. S5c**), the dot product was calculated between the choice axis and the readout axis.

All angles were calculated after first reducing neural activity to the top 10 PCs (PCs calculated from delay period activity; indicated in **Fig. S8b**); for (3), the output weights **W_1_** were projected to these top PCs and unit-normalized. For comparison to minimally structured activity, randomly generated vectors (unit-normalized) were substituted for each axis vector, from which angles were re-calculated 1000 times.

### Neural encoding

To clarify how internal activity in RNNs generated specific behaviors (e.g. RT patterns), we examined activity projections (encodings) along two behaviorally relevant activity axes: (i) the readout axis and (ii) choice axis. For (i), the encoding was defined simply as *z*_1_ (the dot product of activity and the readout axis, with a constant bias term added; see Model architecture, RNN); for (ii), the encoding was defined as the dot product of activity with the choice axis, and was calculated in the top 10 PCs of delay period activity across trial types. All activity vectors were taken under noise-free simulations (*η_i_* set to 0).

To evaluate how end items (item A or G) were encoded during the delay period, we defined an index that quantifies this difference (encoding index) as the ratio of the magnitude of projected activity from the end vs. start of the delay (last vs. first time step, respectively), averaged for A and G trials. This index was calculated for activity projected along either the readout axis (presented in **Fig. 7**) or the choice axis (presented in **Fig. S8c**), which yielded similar results.

### Animal behavioral data

For additional comparison to models (**Fig. S4**), we present data from a behavioral study of traditional TI in Rhesus macaques (originally presented in [53, 161]). The research was approved by the Institutional Animal Care and Use Committee of Columbia University (AAAI1488).

### Human study data

#### Participants

The human research was approved by the Institutional Review Board (IRB) at Columbia University through Columbia IRB Protocol #AAAI1488. All participants provided informed consent for their participation in the experiment. A total of 392 Amazon Mechanical Turk participants took part in the experiment. To promote task engagement, subjects were given a bonus for correct responses (up to $5; availability of bonus stated in advance). As clarified below, a subset of subjects were not included in analyses because they failed to meet pre-established criteria for online studies; subjects were additionally restriected those based in the US within the age range of 18–36 and with an approval rate of above 90%.

#### Materials

The item panel consisted of 7 images of fractals, assigned to items A through G randomly for each subject (**Fig. 8a**). The images were 250 × 250 pixel size and individually presented on top of a white square with a light grey frame (RGB: 200, 200, 200; square size was 265 × 265 pixel size). These stimuli were in turn presented on a grey background (RGB: 128, 128, 128).

#### Task comprehension

Prior to the task, subjects were informed of the basic task format (i.e. presentation of two images, selection of item 1 vs. item 2 images via key presses, ability to choose once item 2 appears and during the choice period) and given a comprehension quiz. Once subjects answered all quiz questions correctly, the task began. Participants were not informed of the underlying relationship between items, or of any underlying difference in trials as the task progressed, specified as follows.

#### Trial structure and responses

Following the trial structure in neural models (**Fig. S1**), each trial consisted of three periods: rest (1 sec), delay (1 sec), and choice (up to 2 sec) (a schematic of trial structure in **Fig. 8b**). The trial began with the presentation of item 1 for 250 ms. A fixation cross (“+”) was then presented for 2 sec, followed by the presentation of item 2 for 250 ms. Next, a choice screen was shown wherein the phrase “make a choice” was presented for up to 750 ms. Participants were instructed to choose one of the items by pressing either the “D” or “K” keys (randomly assigned for each participant to item 1 and 2, or *vice versa*). Participants were instructed in advance that they could respond (make a choice) from the onset of the second item until the end of the choice screen. If participants made a choice during the presentation of item 2, item 2 was nonetheless shown for its full duration (250 ms), but the choice screen was not subsequently shown in that trial. A response made during the choice screen terminated the choice period/screen; immediately thereafter, a screen indicating the outcome of their choice (green check: correct response; red cross: incorrect response) for 1 sec. If participants did not respond during the allotted time, they were shown a screen displaying “too slow” for 1 sec, and were not shown the outcome screen; these trials were treated as incorrect trials in calculating performance (% correct of all trials). In trials in which subjects responded, the response time (RT) was measured as the time elapsed from the onset of presentation of item 2. The next trial automatically began after a rest period of 1 sec.

#### Task phases and trial blocks

The task consisted of two phases, the training phase and testing phase. The training phase consisted of 144 trials divided into 3 blocks of 48 trials each, with these trials consisting of 4 repetitions of the 12 training trial types (A vs. B, B. vs. C, etc.), presented in a subject-specific random order in each block. The testing phase consisted of 252 trials divided into 6 blocks of 42 trials each, with these trials consisting of all training trial types (12 trials) and all testing trial types (30 trials), presented in a subject-specific randomized order in each block. Upon completion of testing, subjects were asked by questionnaire to describe how they performed the task. The entire duration of the task was ~40 min on average. It is also worth noting that the degree of training is relatively low in the present paradigm, particularly in comparison to TI task paradigms involving thousands of training trials.

#### Inclusion criteria

To ensure that subjects were attending to the task, the following inclusion criteria were imposed: (i) fewer than 20 trials in which no response was given; (ii) fewer than 20 events where participants were browsing a different window in any experimental phase (blur-focus events detected using jsPsych library [162]), and (iii) fewer than 10 failed attempts to pass the comprehension quiz. Further, to ensure that subjects had the minimal degree of task proficiency prerequisite to testing generalization, a criterion of (iv) above chance performance (>50%) on training trials (i.e. A vs. B, B vs. C, etc.) in the final training block (block 3, **Fig. 8c-d**) was imposed. In an initial evaluation of inference in the delay TI paradigm, in place of (iv) a more stringent criterion of >80% performance on training trials in the final training block was imposed, similar to previous studies that initially establish TI task paradigms [51, 115]. In all main analyses (**Fig. 8** and **Fig. S8**), only subjects meeting criteria (i)-(iv) were included.

### Statistical tests

All statistical tests were non-parametric and two-sided.

## Supplementary Figures and Diagram

**Fig. S1.**
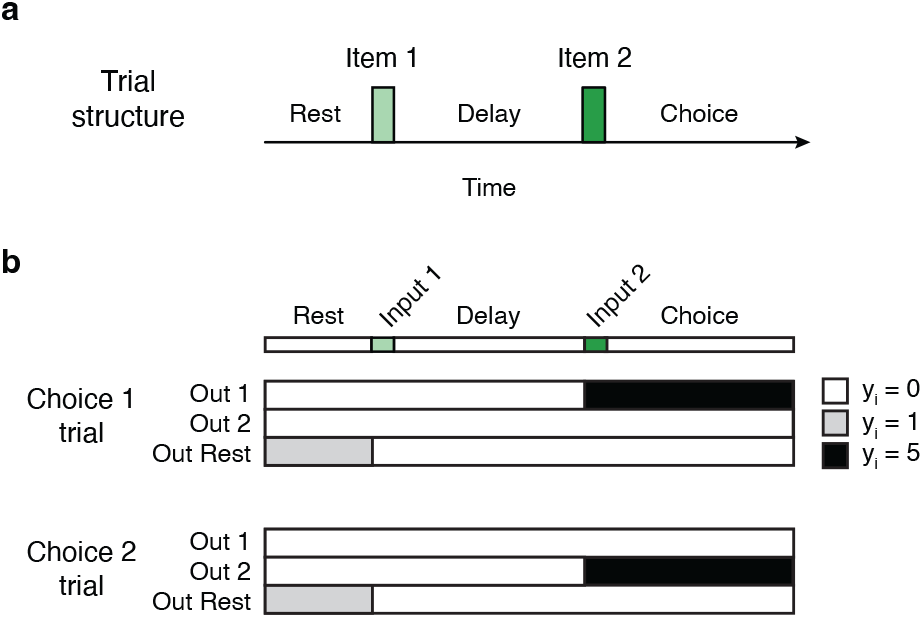
Delay TI: transitive inference (TI) with a requirement for working memory. **a**, Trial structure. Each trial consists of three periods: rest, delay, and choice. Note that subjects respond on the basis of item order: if the correct response in trial type X vs. Y (item 1: X, item 2: Y) is choice 1, then the correct response in trial type Y vs. X (item 1: Y, item 2: X) is choice 2. **b**, Target values of RNNs output units (*z_i_*(*t, m*), where t is time and m is trial type; see Methods).

**Fig. S2.**
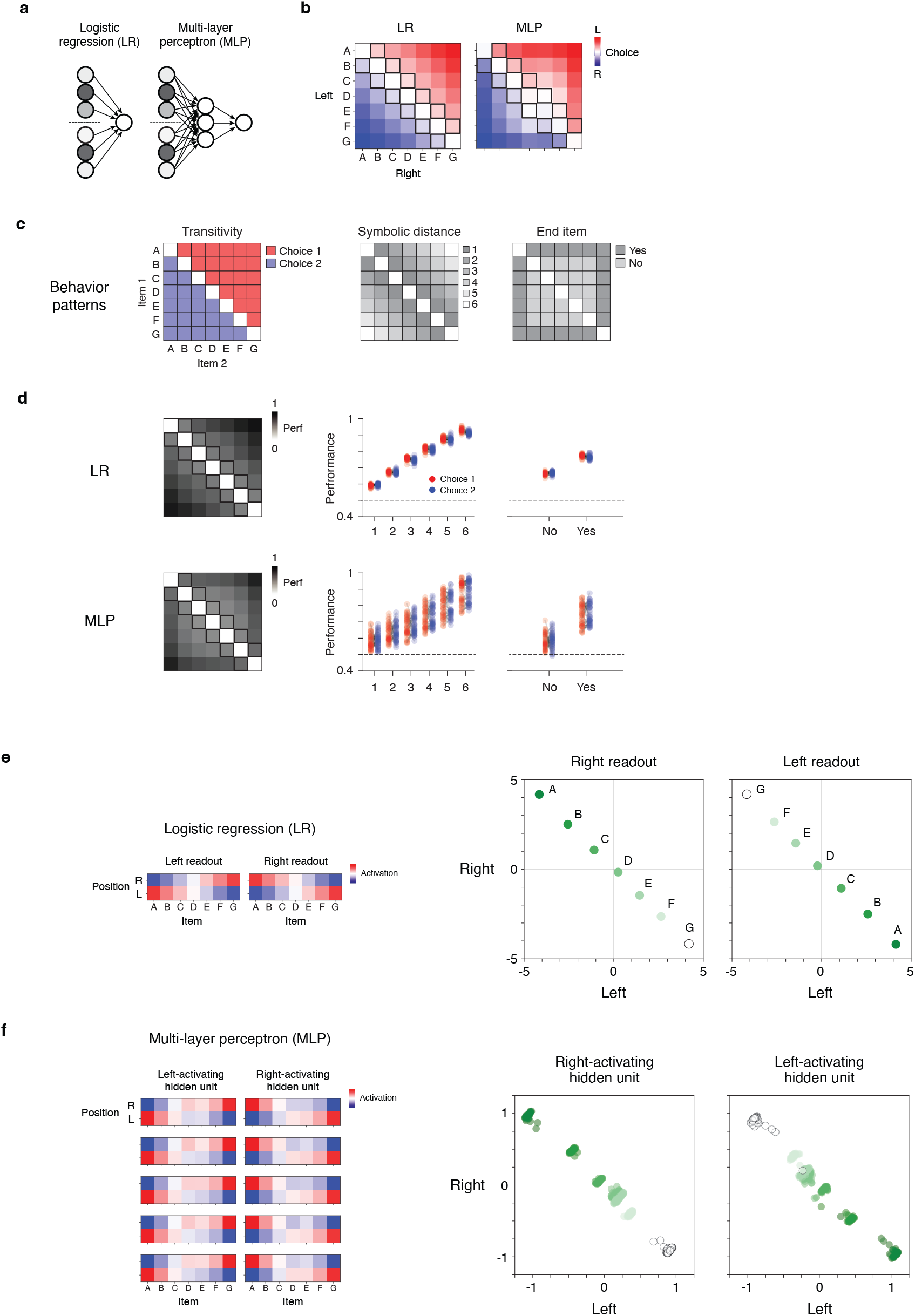
Traditional TI in feedforward models. **a**, Schematic of feedforward model architecture (see Methods). **b**, Example LR and MLP model instances that perform traditional TI (i.e. no explicit delay between items, with choice made on basis of position (left vs. right); Fig. 1b). **c**, Schematic of behavior patterns. **d**, Behavior of feedforward models (n = 100 instances / model). All plots show average performance (proportion correct, averaged across 500 simulations of every trial type). Column 1: Averages across model instances by trial type. Columns 2-4: Averages across trials for each model instance by trial type. Trial types follow that defined for each behavioral pattern in panel c (column 2: symbolic distance; column 3: end item), in addition to distinguishing between choice 1 vs. choice 2 trial types (red vs. blue, respectively; diagrammed in panel c, transitivity). **e** and **f**, feedforward models express a ‘subtractive’ solution to TI. **e**, Analysis of an example LR. At left, activation of readout ‘units’ (see Methods) as a function of input position (y-axis) and rank (x-axis). At right, relationship between position of inputs and readout unit activation. Note that activations by item position (left vs. right) were sign-inverted versions of each other. **f**, Analysis of an example MLP. At left, activation of hidden units as a function of input position (y-axis) and rank (x-axis). At right, relationship between position of inputs and unit activation, plotted for all hidden units (N=100 tanh units). Note that activations by item position (left vs. right) were approximately sign-inverted versions of each other, akin to the LR model.

**Fig. S3.**
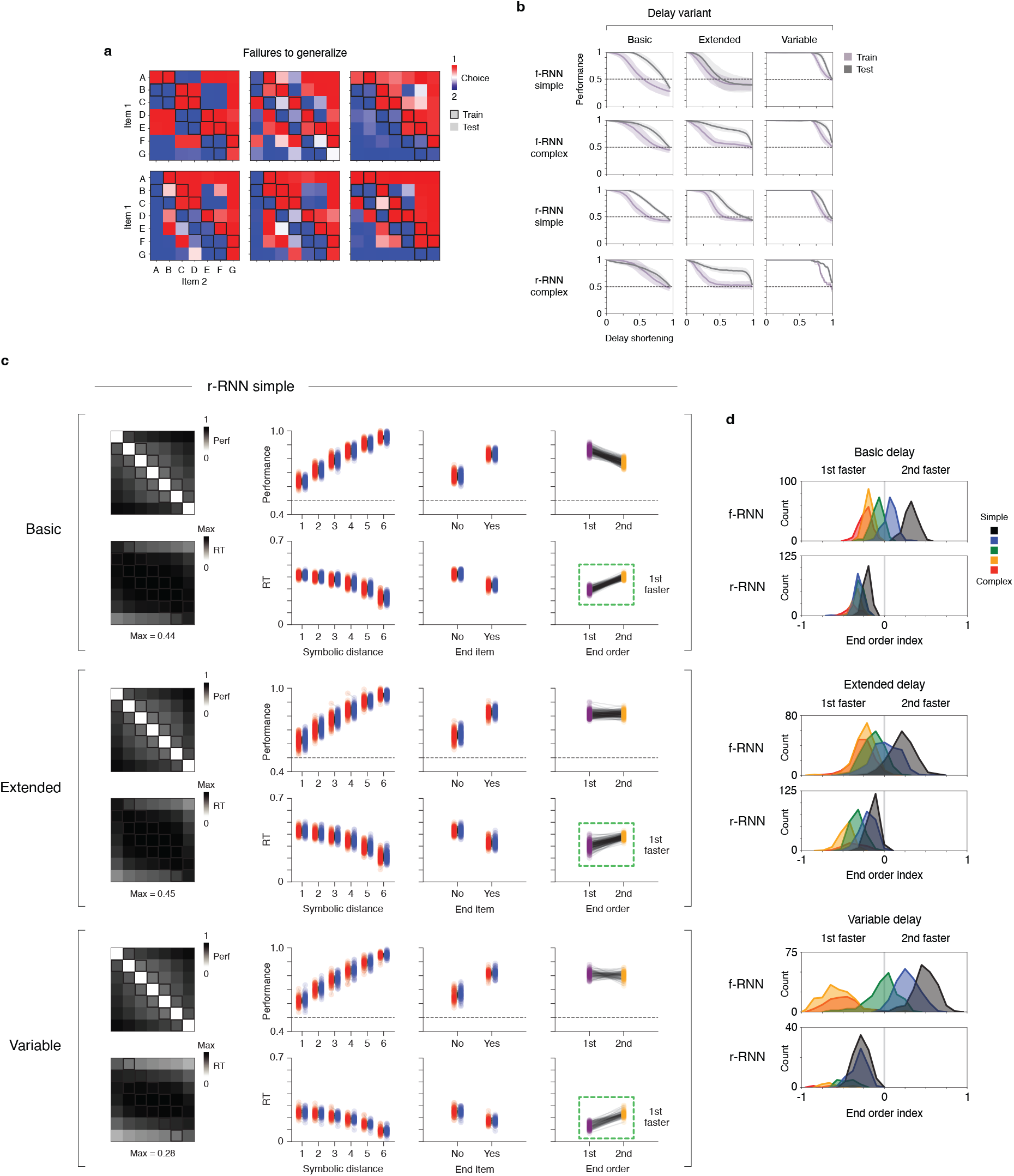
Delay TI in additional RNNs. **a**, Six example RNNs that responded correctly in training trials, but failed to generalize. Plotted are network outputs by trial type (compare to Fig. 2, top row, plotting conventions shared). **b**, Performance (proportion correct) as a function of delay length. RNNs were trained on three delay variants: basic, extended, and variable (see Methods, Model input). Performance was measured when trials were shortened relative to the fixed (basic and extended) or maximal (variable) delay length, and performance was measured separately for training (purple) vs. test (grey) trial types. Plots show averages (dark traces) ± s.d. (shaded regions) across model instances (see **Table 4** for model counts). **c**, Behavioral patterns in simple-regime r-RNNs across delay variants. Plotting conventions are the same as in Fig. 3. **d**, End order behavior across delay variants (see Methods). Plotting conventions are the same as in Fig. 4, with x-axis range (−1 to +1) made equivalent across plots to aid comparison.

**Fig. S4.**
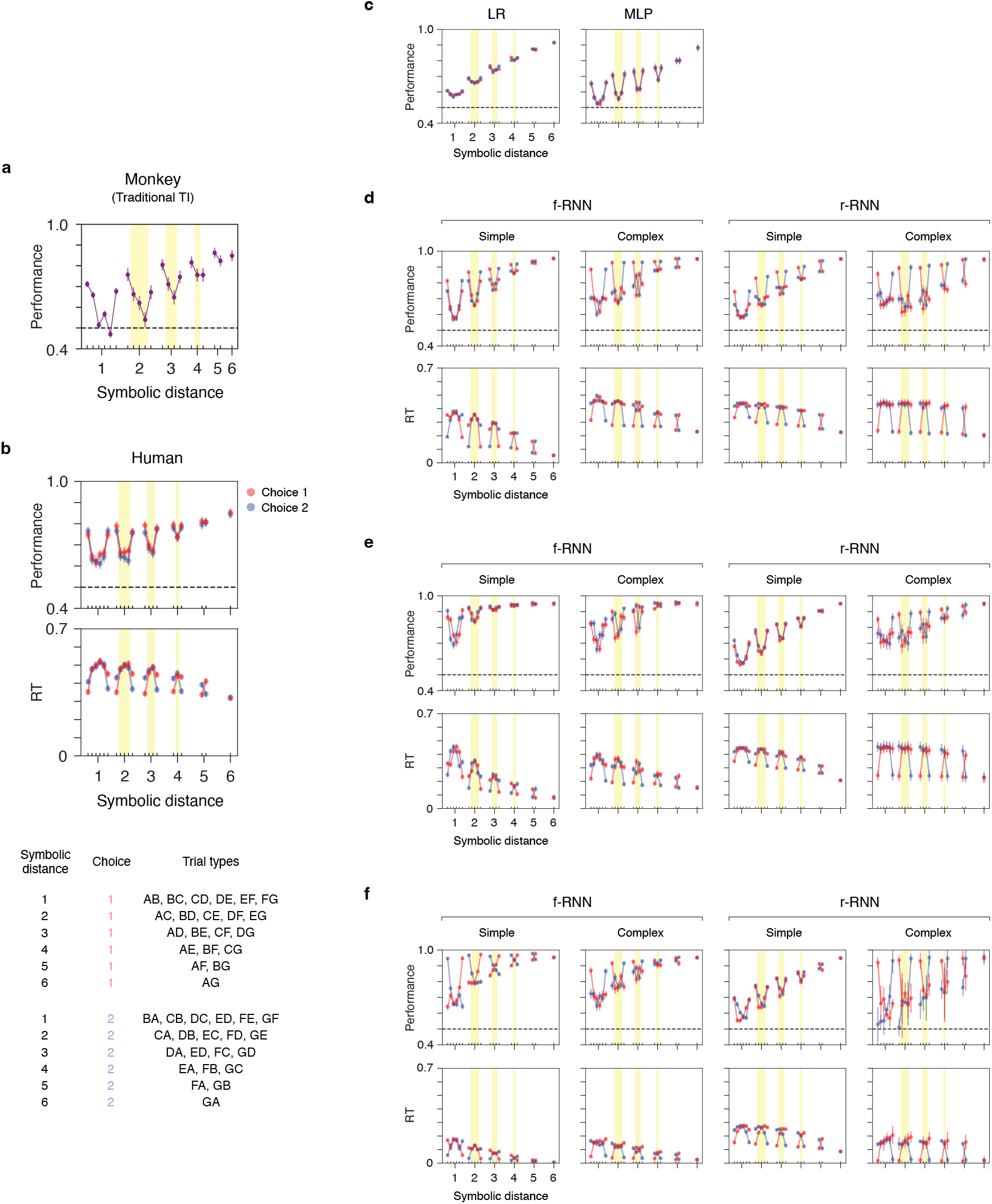
RNNs show TI behavior similar to that of living subjects. Comparison of behavior across trial types: living subjects and models. The behavioral data plotted are similar to that of Fig. 3, but here more explicitly show differences across trial types. **a**, Monkey performance in traditional TI (items presented simultaneously, Fig. 1c) by trial type. Trial types defined solely by rank of items and not order (by symbolic distance, 1: AB, BC, CD, DE, EF, FG; 2: AC, BD, CE, DF, EG; 3: AD, BE, CF, DG; 4: AE, BF, CG; 5: AF, BG; 6: AG). Originally reported in [53, 161]. **b-f**, Human and RNN performance and response times (RTs) in delay TI by trial type (n=292 human subjects; see **Table 4** for numbers of RNN instances). In delay TI, trial types depend additionally on order (AB, BA, BC, CB, etc.). Plotted are average performance (top row) and RTs (bottom row; not available in feedforward models); trial types in each plot, from left to right for each symbolic distance, are at the bottom of panel **b**, with the distinction of choice 1 vs. 2 trials (red and blue, respectively). Highlighted in each plot are ‘critical’ trial types (testing trials that do not contain end items (A or G); yellow zones). **c**, Feedforward models (LR and MLP; n=100 instances / model type). **d-f**, RNNs (**d**, **e**, and **f** corresponding to three delay variants: basic delay, extended delay, and variable delay; columns: f-RNN/r-RNN and simple/complex (simple (high) and complex (high))). Error bars are ± 1 s.e.m. (monkey and human subjects) and ± 2 s.e.m. (all models).

**Fig. S5.**
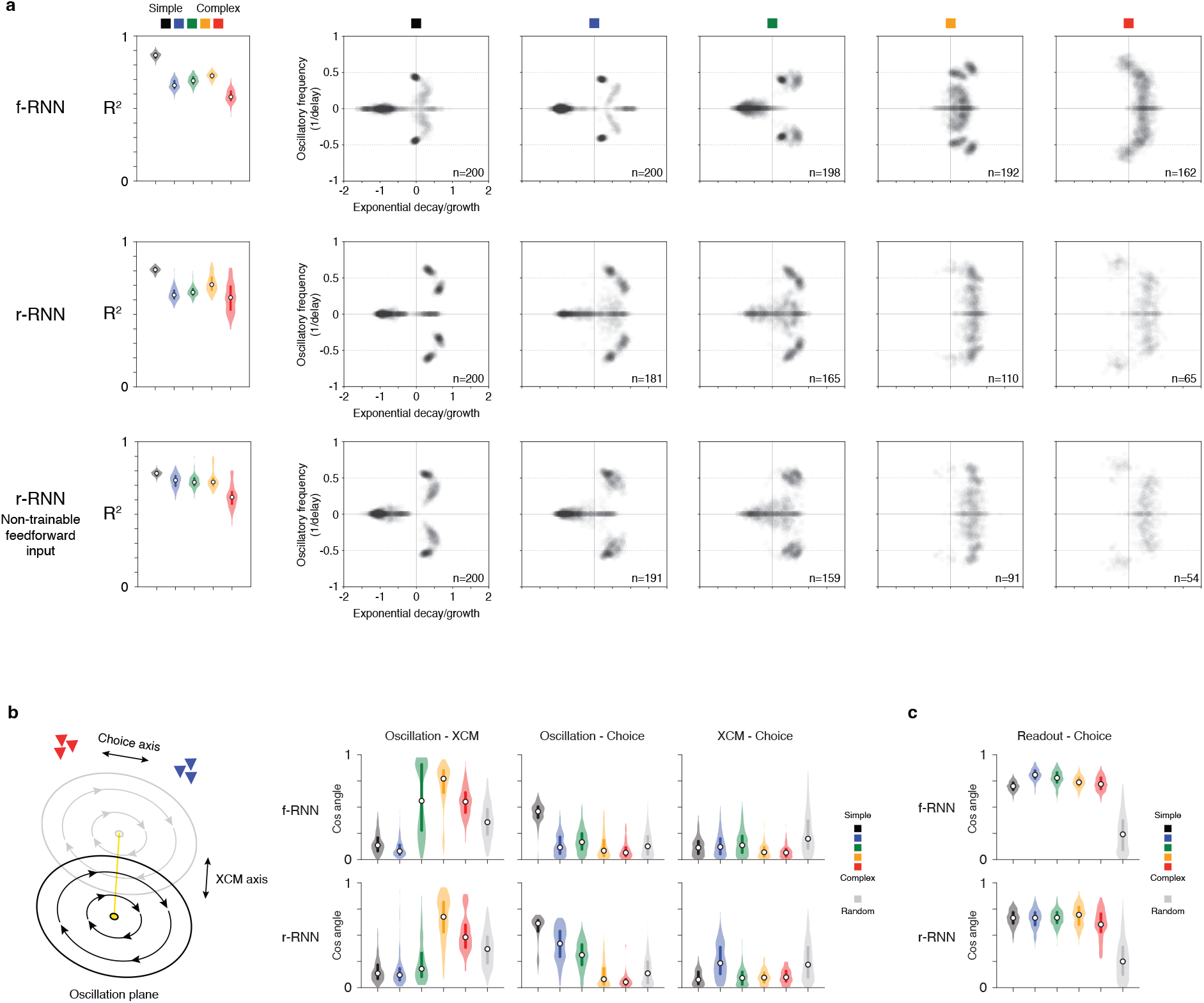
RNNs performing delay TI: linear dynamics and geometric alignments. **a**, RNN activity during the delay period was fit to a linear dynamics model (least-squares). Rows show results for three RNN variants differing by learnable connectivity (f-RNN: fully-trainable RNN, r-RNN: recurrent-trainable RNN, and last row corresponds to r-RNN for which only feedforward inputs (not outputs) were not trainable). Column 1: *R*^2^ values of the fit. Training regime variants plotted by color. Columns 2-5: eigenvalue spectra (grey points; calculated for each instance using top 10 PCs). Each column corresponds to a different RNN variant (simple to complex; n=65-200 instances). Note that spectra shown in Fig. 5c corresponds to two of the spectra here combined (f-RNN simple (high) (black, second column) and simple (low) (blue, third column)), and that spectra shown in **Fig. S6c** is the same as that shown in the third row, second column. **b,** Alignment between activity axes. **Left**, Schematic of activity geometry expressed in RNNs (compare to **figS. 5a** and **S6a**). The oscillation plane refers to the oscillatory mode associated with transitive comparison (if expressed in network activity). Note that simple-regime RNNs (**figS. 5** and **S6**) make two specific predictions: (1) the oscillation plane should be orthogonal to the XCM axis (cosine angle: ~0), and (2) the oscillation plane should be aligned to the choice axis (cosine angle: larger than random). All measures were inferred from neural activity under noiseless conditions. XCM: cross-condition mean. **Right**, Quantification of alignment of the activity geometry shown in panel b (cos angle: 1 (fully aligned), 0 (orthogonal)). For additional comparison, values obtained between random activity vectors are shown (light grey). The quantification clarifies the predictions of simple-regime RNNs stated above; for summary of predictions, see **Table 3**. **c**, Alignment of readout axis and choice axis. Note that the axes are consistently aligned across all RNNs. We also observed that for nearly all RNNs performing TI (98% of all instances), output units showed full separation between choice 1 vs. 2 (across all trial types) within the first ~10% of the choice period (equivalent to ~10% of the delay period).

**Fig. S6.**
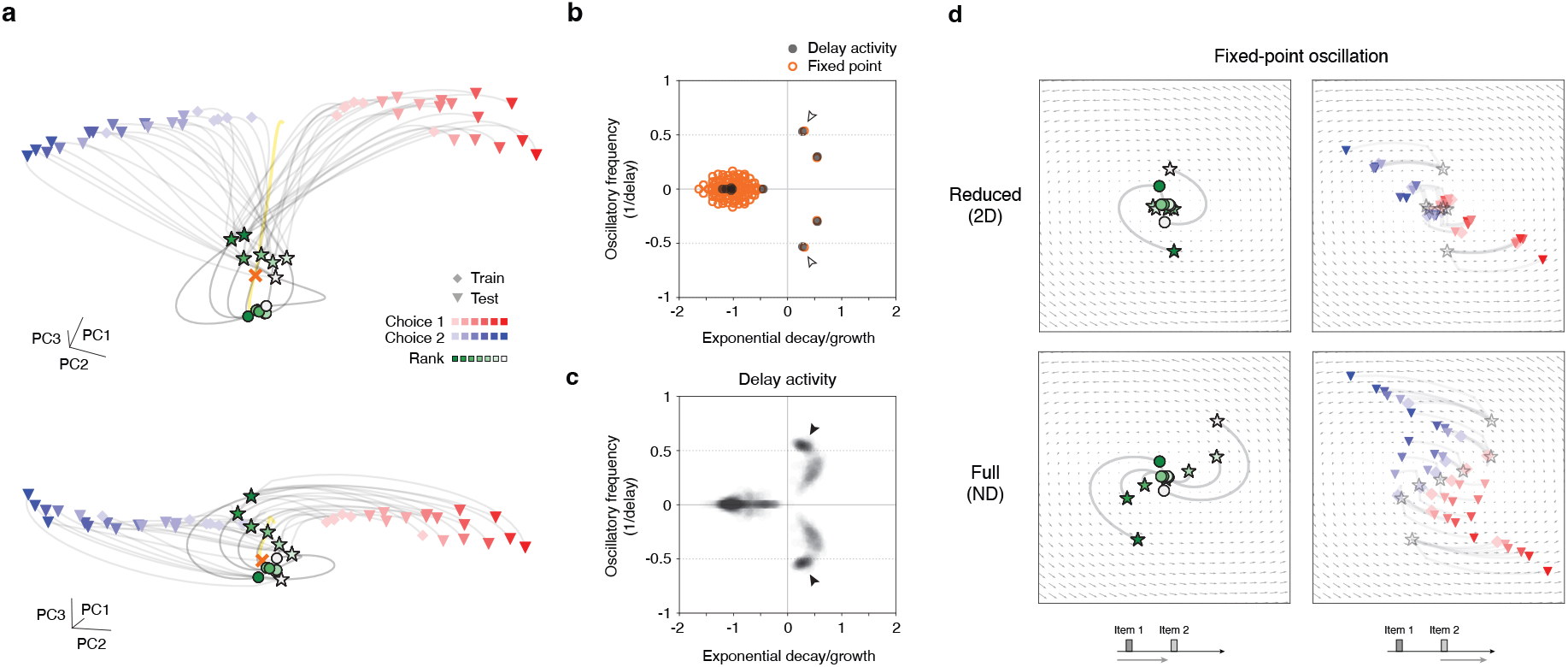
A single oscillation enables TI in r-RNNs. **a**, Population activity trajectories in an RNN (simple-regime r-RNN) that performs TI. Top and bottom plots show two different views. Shown are trajectories from all 42 trial types (Fig. 1b). To clarify the operation of the network, three trial times are highlighted as follows: (i) presentation of item 1 (green circles; shade indicating item rank: A (dark green) to G (white)), (ii) the last time point of the delay period (green stars; same color convention), (iii) last time point of the trial (red/blue symbols; red: choice 1 trials, blue: choice 2 trials, light to dark shading indicating symbolic distance (1 to 6); diamonds: training trials, triangles: test trials). Also shown: cross-condition mean (XCM; the average trajectory across all trial types) (yellow line) and fixed point (FP) (orange cross). The FP was located near trajectories during the delay period (‘early-trial’ FP, compare to Fig. 6a). Note the oscillatory evolution of trajectories in the delay period (circles to stars) despite the absence of a linearly arranged rank-ordered activity upon presentation of item 1 (green circles; compare to Fig. 6a). **b**, Linear dynamics of RNN in panel a. Two eigenvalue spectra of the RNN are plotted: first, the spectrum calculated from delay-period neural activity (black points; inferred via least-squares linear fit, *R*^2^ = 0.78) and second, the spectrum from linearization of the network with respect to the early-trial FP (orange circles; FP shown as orange cross in panel a). **c**, Linear dynamics of simple-regime f-RNNs (n=200 instances, simple (high) regime). Eigenvalue spectra of delay-period neural activity (grey translucent points; inferred via least-squares linear fit, *R*^2^ ~ 0.8 across the 200 instances, see **Fig. S6**). Note the density of oscillatory modes with frequency ~0.5 cycles / delay (filled arrowhead; compare to Fig. 5c). **d**, Activity trajectories in the oscillatory mode of the linearized RNN. The oscillatory mode is that of the linearization of the early-trial FP (open arrowheads in panel b). Two sets of trajectories are plotted (rows): one where all activity (initial conditions and inputs) has been reduced to the 2 dimensions of the oscillatory mode (top row, reduced) and one where the full dimensionality (N-D; here N=100) of initial conditions and inputs have been retained (bottom row, full). In both cases, the dynamics are governed solely by the oscillatory mode (see Methods), though only in the reduced (2D) case do trajectories strictly follow the flow field vectors. To clarify the progression of activity, trajectories are plotted in two stages of the task trial (left and right columns; schematic of the stages at the bottom of each panel): early trial (left) and late trial (right). To clarify how the activity evolves, specific trial times are highlighted as follows: (i) presentation of item 1 (green circles; shade corresponding to item rank: A (dark green) to G (white)), (ii) the last time point of the delay period (green stars; same color convention), (iii) a quarter of the time period following presentation of item 2 (i.e. choice period, see **Fig. S1**; red/blue symbols; red: choice 1 trials, blue: choice 2 trials; diamonds: training trials, triangles: test trials). Note that separation (linear separability) of choice 1 vs. 2 trials (red vs. blue symbols) does not occur in the reduced-activity case, but does occur in the full-activity case.

**Fig. S7.**
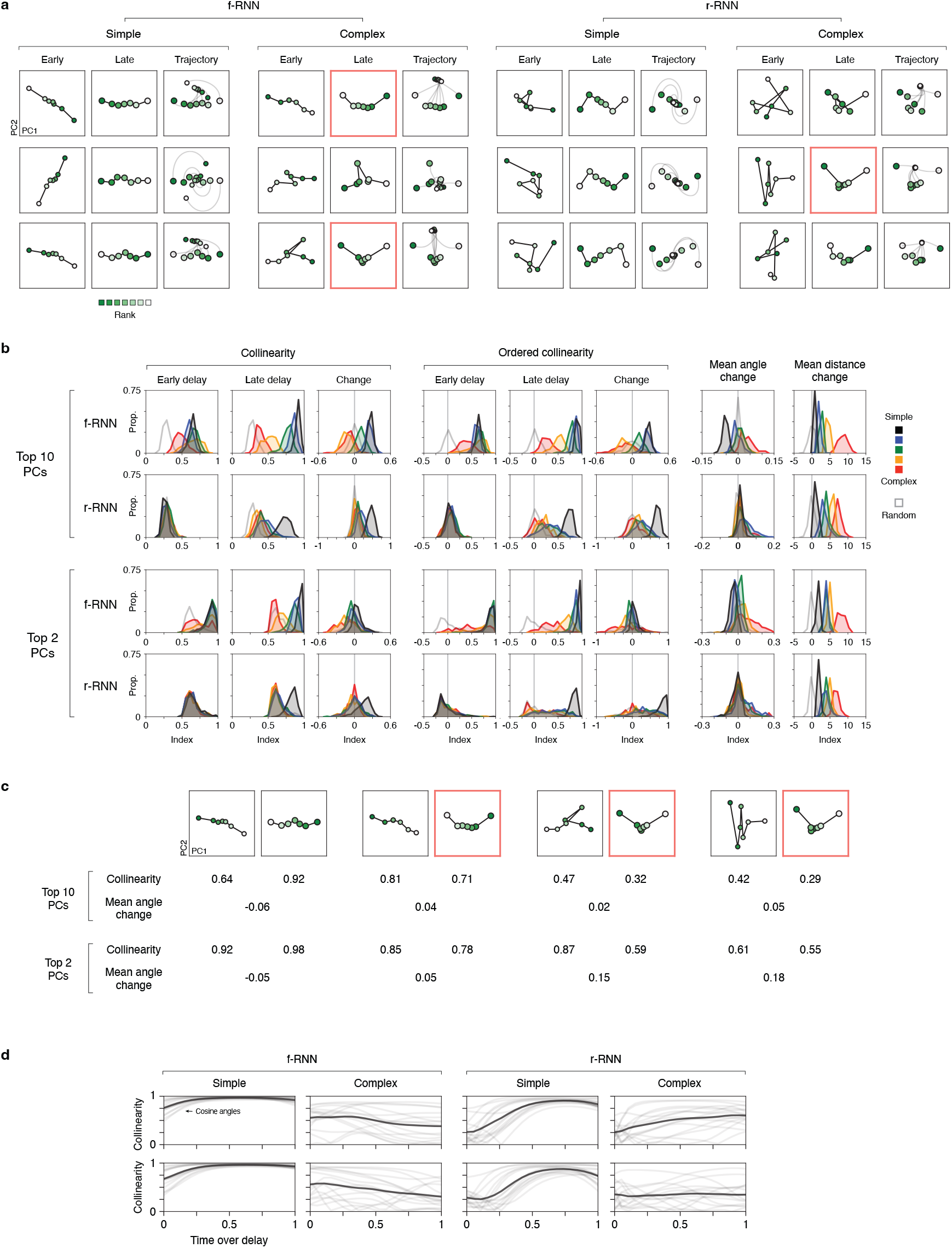
RNNs performing delay TI: activity geometry during the delay. **a,** Delay period population activity in 12 additional example RNNs. Plotting conventions follow that of Fig. 6a. In instances of complex- regime RNNs, a “U” activity geometry was expressed by the end of the delay period (late delay; highlighted with red box). In the instances shown, the mean angle change index values were 0.04 (top row, f-RNN complex), 0.02 (bottom row, f-RNN complex), and 0.05 (middle row, r-RNN complex). **b**, Histograms of geometric index values across RNNs. Plotting conventions follow those of Fig. 6b**-c**, with the addition of ordered collinearity (see Methods) and with the same analysis carried out in the top 2 PCs (at bottom). All plots show histograms of instances for each RNN variant (n=65-200 instances / variant; see **Table 1**), in addition to randomly generated data (open grey histograms). **c**, Two geometric indices for four example RNNs. Each example (column sections) is from panel a (f-RNN simple, f-RNN complex, f-RNN complex, r-RNN complex). At top, the delay period population activity (PC1 and PC2; early delay (left) and late delay (right)) is shown. At bottom, geometric index values are shown, calculated in the top 10 and top 2 PCs. Note that the second through fourth examples show “V” shaped geometry in late delay, and further have positive mean angle change values. **d**, Collinearity over the course of the delay period in eight example RNNs (two examples / variant; variant indicated above). In each plot, two measures are plotted: the collinearity index (black lines; schematized in Fig. 6b) and individual cosine angles between trial types (grey lines; e.g. A vs. B, B vs. C activity states). The collinearity index is the average across cosine angles.

**Fig. S8.**
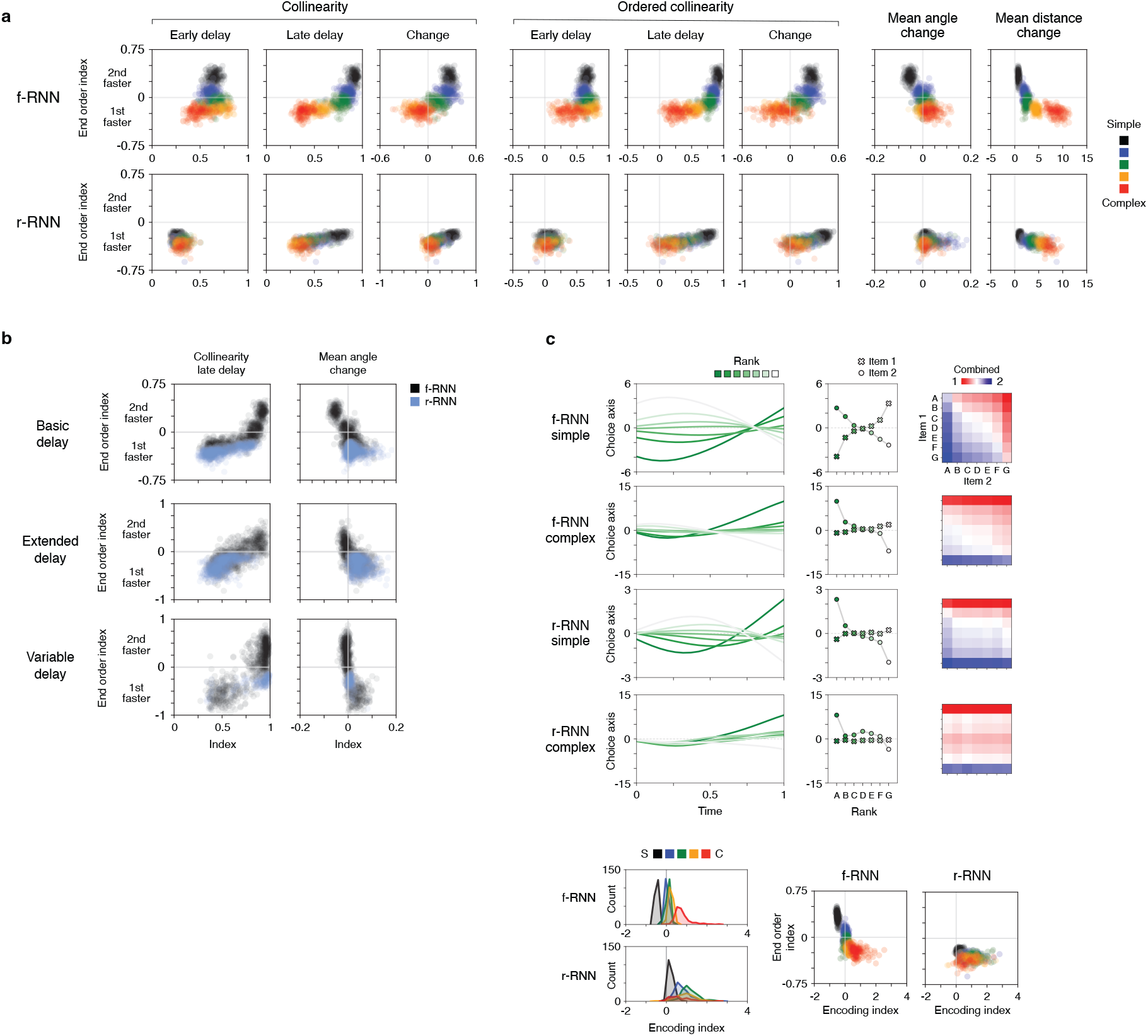
RNNs performing delay TI: activity geometry and encoding strategy predict behavior. **a**, End order behavior vs. activity geometry across RNN variants. Behavior (y-axis) is the end order pattern (figS. 3 and 4; quantified by the end order index; see Methods), for which RNNs show alternative versions (1st vs. 2nd-faster; >0 and <0 index values, respectively). Activity geometry (x-axis) correspond to the patterns schematized and quantified in **figS. 6** and **S7**. **b**, End order behavior vs. activity geometry across all RNNs in the present study. Each row corresponds to a different delay variant (see Methods for detials). Plots contain the same data as in panel a, but do not show training regime. Note that for mean angle change, alternative behaviors (index values <0 vs. >0) correspond to qualitatively different geometries (>0 vs. <0). **c**, Alternative encoding strategies in RNNs. Example networks (upper rows) and plotting conventions are the same as in Fig. 7, with the difference that activity projections were on the choice axis (rather than readout axis).

**Fig. S9.**
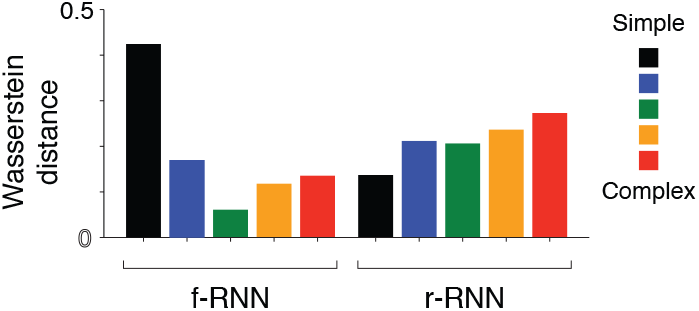
End order effect: RNNs vs. human data. To compare model behavior to that of human subjects, the Wasserstein distance (earth mover’s distance) was calculated between the end-order index values across instances for each RNN variant (Fig. 4b) to index values across human subjects (Fig. 8g).

**Supplemental diagram.**
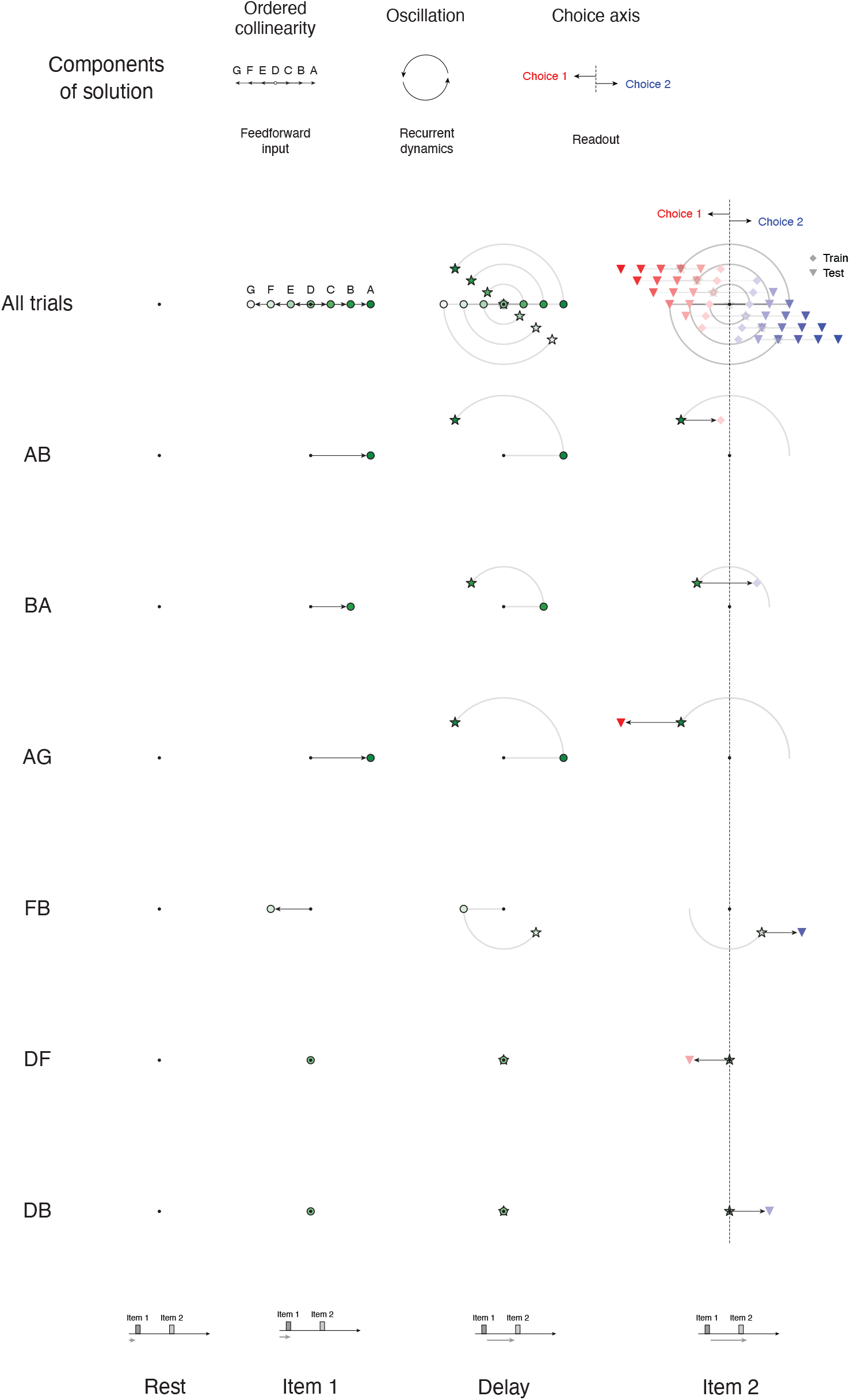
The “subtractive” solution to delay TI. Diagrams presenting the solution in greater detail (compare to Fig. 5). Top, diagram of each of the population-level components comprising the solution (top: the specific form of the component; bottom: the network implementation). Bottom, activity trajectories across trial periods (columns; diagram of each period at bottom) and across different trial types (rows; top row: all trials; single trial types in rows below). Trajectories were generated by simulating a 2D linear dynamical system defined by an oscillation of frequency ~0.5 cycles / delay, with initial condition at the origin and input vectors encoding task items (A, B, C, etc.) in ordered collinear arrangement in state space. Trial-based input (item 1 - delay - item 2, see **Fig. S1a**) was applied to the system.

